# Inherited retinal degeneration: T-type voltage-gated channels, Na^+^/Ca^2+^-exchanger and calpain-2 promote photoreceptor cell death

**DOI:** 10.1101/2023.07.16.549200

**Authors:** Jie Yan, Lan Wang, Qian-Lu Yang, Qian-Xi Yang, Xinyi He, Yujie Dong, Zhulin Hu, Kangwei Jiao, François Paquet-Durand

## Abstract

Inherited retinal degeneration (IRD) refers to a group of untreatable blinding diseases characterized by a progressive loss of photoreceptors. IRD pathology is often linked to an excessive activation of cyclic nucleotide-gated channels (CNGC) leading to Na^+^– and Ca^2+^-influx, subsequent activation of voltage-gated Ca^2+^-channels (VGCC), and further Ca^2+^ influx. However, whether and how exactly intracellular Ca^2+^ overload contributes to photoreceptor degeneration is still controversial.

Here, we used whole-retina and single-cell RNA-sequencing to compare gene expression between the *rd1* mouse model for IRD and wild-type (*wt*) mice. Differentially expressed genes were linked to several Ca^2+^–signalling related pathways. To explore this further, organotypic retinal explant cultures derived from *rd1* and *wt* mice were treated with the intracellular Ca^2+^-chelator BAPTA-AM and with inhibitors for different Ca^2+^-permeable channels, including CNGC, L-type VGCC, T-type VGCC, Ca^2+^-release-activated channel (CRAC), and Na^+^/Ca^2+^ exchanger (NCX). Moreover, we employed the compound NA-184 to selectively inhibit the Ca^2+^-dependent protease calpain-2. The overall activity of poly(ADP-ribose) polymerases (PARPs), sirtuin-type histone-deacetylases, calpains, as well as the activation of calpain-1, and –2 were analysed *in situ* on retinal tissue sections. Cell viability was assessed *via* the TUNEL assay.

While *rd1* photoreceptor cell death was reduced by BAPTA-AM, the effects of Ca^2+^-channel blockers were ambiguous, with T-type VGCC and NCX inhibition showing protection, while blocking CNGC and CRAC was detrimental. Activity of calpains and PARPs generally followed similar trends as cell death. Remarkably, sirtuin activity and calpain-1 activation was associated with photoreceptor protection, while calpain-2 activity was linked to degeneration. Accordingly, the calpain-2 inhibitor NA-184 protected *rd1* photoreceptors.

Together, these results indicate that Ca^2+^ overload in *rd1* photoreceptors may be triggered by T-type VGCC in conjunction with NCX. High Ca^2+^-levels likely suppress the protective activity of calpain-1 and promote neurodegeneration via activation of calpain-2. Our study details the complexity of Ca^2+^-signalling in photoreceptors and emphasizes the importance of identifying and targeting degenerative processes to achieve a therapeutic benefit for IRD.

## INTRODUCTION

Inherited retinal degeneration (IRD) is the hallmark of a large group of genetically heterogeneous blinding diseases [1]. The most common disease within the IRD group is *retinitis pigmentosa* (RP), with a prevalence of approximately 1:4000, affecting more than two million patients worldwide [2]. RP patients initially experience reduced night vision due to primary degeneration of rod photoreceptors, and then suffer from a progressive visual field loss due to secondary degeneration of cone photoreceptors [2]. The second messenger cyclic-guanosine-monophosphate (cGMP) has been found to play a central role in the pathobiology of many genetically distinct types of IRD [3], and may be directly or indirectly associated with the activity of histone deacetylases (HDACs), poly(ADP-ribose) polymerases (PARPs), cyclic nucleotide-gated channels (CNGCs), and calpain-type proteases [3–5].

Phototransduction in photoreceptors critically depends on Ca^2+^– and cGMP-signalling. cGMP is produced by guanylyl cyclase, the activity of which is inhibited by Ca^2+^ [5]. In darkness, cGMP opens CNGC causing influx of Na^+^ and Ca^2+^. Together, Ca^2+^ and cGMP form a feedback loop which controls the levels of both second messengers. CNGC-mediated ion influx is countered by the Na^+^/Ca^2+^/K^+^ exchanger (NCKX) and by the ATP-driven Na^+^/K^+^ exchanger (NKX) [5]. As a result, in the dark, a photoreceptor cell is depolarised at approximately –40 mV [6, 7]. The consequent activation of voltage-gated Ca^2+^-channels (VGCC) mediates further Ca^2+^ influx and synaptic glutamate release [6, 8]. In light, the enzyme phosphodiesterase-6 (PDE6) rapidly hydrolyses cGMP, leading to CNGC closure, Ca^2+^ decrease, and photoreceptor hyperpolarization. Subsequently, VGCC closes, ending synaptic neurotransmitter release [5]. In IRD, a mutation-induced cGMP accumulation is likely to over-activate CNGC and to produce an abnormal influx of Na^+^ and Ca^2+^ into photoreceptors.

The *rd1* mouse is a naturally occurring IRD mouse model first described by Keeler in the early 1920s [9]. It is characterized by early onset, rapid retinal degeneration due to a mutation in the rod-photoreceptor-specific *Pde6b* gene [10]. This causes dysfunction of cGMP-phosphodiesterase-6 (PDE6) and subsequent accumulation of cGMP in rods. This in turn results in primary rod degeneration, starting at postnatal day 9 (P9), with a peak of cell death at about P13, and with essentially all rods lost by P21. This is followed by secondary cone photoreceptor cell loss occurring between P18 to P60 [11, 12]. The primary *rd1* degeneration has often been connected to an excessive Ca^2+^-influx, notably through cGMP-dependent CNGCs [13]. Excessive Ca^2+^ is thought to promote activity of Ca^2+^-dependent calpain-type proteases and *rd1* photoreceptor death [14]. Accordingly, many studies over the past decades have focused on the role of Ca^2+^ in IRD, albeit without conclusive results until today [15]. While some studies suggested that Ca^2+^– channel blockers or genetic ablation of Ca^2+^-permeable channels could preserve photoreceptors [16, 17], others argued that inhibition of Ca^2+^-permeable channels either does not reduce retinal cell death or may even promote degeneration [18, 19]. Overall, these conflicting results raise the question whether *rd1* rod cell death is linked to Ca^2+^ overload, and, if so, which Ca^2+^-permeable channels might be responsible.

A candidate for a Ca^2+^-permeable channel causing *rd1* pathology is VGCC. Functional VGCCs are composed of pore-forming α_1_ subunit proteins, encoded by *CACNA1x* genes, of which there are 10 isoforms in the mammalian genome [20]. In the case of Ca_V_1.1 – 1.4 channels (known as L-type channels), these are encoded by *CACNA1S*, *-C*, *-D* and *-F*, respectively. The Ca_V_2.1 – 2.3 channels (termed P/Q-, N– and R-type) are encoded by *CACNA1A*, *-B* and *-E*, accordingly. The T-type Ca_V_3.1 – 3.3 channels are encoded by *CACNA1G*, *-H* and *-I* [20, 21]. The accessory α_2_δ and β subunits are important for channel folding, subsequent transport to the cell surface, and their integration into specific domains in polarized cells such as neurons. Both Ca_V_1 and Ca_V_2 classes of channels form a heteromeric complex, co-assembling with one of four β subunits (encoded by *CACNB1-4*), and one of four α_2_δ subunits (encoded by *CACNA2D1-4*) [20].

Changes in membrane potential triggered by either CNGC or VGCC activity could affect the activity of the Na^+^/Ca^2+^ exchanger (NCX). This is a bi-directional regulator of cytosolic Ca^2+^, which in forward mode transports Ca^2+^ out of cells; however, in reverse mode NCX may import extracellular Ca^2+^ [22]. As NCX does not require ATP for ion transport, the direction of Ca^2+^ movement through the channel depends on the net electrochemical gradients for Na^+^ and Ca^2+^, such that membrane depolarization augments Ca^2+^ influx [22]. Ion transport by the NCX is electrogenic, with a stoichiometry of three Na^+^ ions exchanged for each Ca^2+^ ion [23]. In mammals, three different NCX genes have been identified: *SLC8A1* encoding NCX1, *SLC8A2* encoding NCX2, and *SLC8A3* encoding NCX3 [24].

Store-operated Ca^2+^ entry (SOCE) is a ubiquitous Ca^2+^ signalling pathway, which is evolutionarily conserved across eukaryotes. SOCE is triggered physiologically when the endoplasmic reticulum (ER) Ca^2+^ stores are depleted. In this situation Ca^2+^-levels are replenished through Ca^2+^-release-activated Ca^2+^-channels (CRAC) [25]. SOCE involves a complex choreography between the plasma membrane (PM) protein “Orai” (encoded by *ORAI1* and *ORAI2*) and the ER-resident Ca^2+^-sensing stromal interaction molecules (Stims, encoded by *STIM1* and *STIM2*). The depletion of ER Ca^2+^ is sensed by STIM1 and its homolog STIM2, causing the opening of Orai channels to drive Ca^2+^ entry into the cell [25].

To try and settle the long-standing controversy on the role of Ca^2+^ in IRD pathogenesis, we performed an initial bio-informatic analysis of differentially expressed genes (DEGs) in *rd1* mouse retina based on whole-retina RNA sequencing (RNA-seq) and single-cell RNA sequencing (scRNA-seq) datasets. We found numerous DEGs to be enriched in Ca^2+^ associated pathways, arguing for an important function of Ca^2+^–signalling. We then used Ca^2+^ chelation and Ca^2+^-permeable channel inhibition in organotypic *rd1* retinal explant cultures to identify the possible sources of high photoreceptor Ca^2+^. To assess the role of Ca^2+^– dependent proteolysis, we also studied the overall activity of calpains, and then focussed on calpain-1 and –2. We found that calpain-2 contributed to photoreceptor cell death, a finding confirmed by the protective effect of the selective calpain-2 inhibitor NA-184. Altogether, we show that Ca^2+^ contributes to *rd1* photoreceptor cell death, regulating, among other things, the activity of PARPs and sirtuins, and that Ca^2+^ overload and photoreceptor degeneration is likely caused T-type VGCC. Surprisingly, inhibition of CNGC or CRAC accelerate retinal degeneration.

## MATERIALS AND METHODS

### Animals

For retinal explant cultures C3H/HeA *Pde6b^rd1/rd1^*animals (*rd1*), their congenic wild-type C3H/HeA *Pde6b*^+/+^ counterparts (*wt*) [26], and B6.129SvJ;C3H/HeA-*CNGB1*^tm^ double-mutant mice (*rd1*Cngb1^-/-^*) were used [17]. The *rd1*Cngb1^-/-^*double mutants were generated by an intercross of *rd1* and *Cngb1^-/-^*[17]. Animals were used regardless of gender. The stock has been maintained by repeated backcrossing over ten generations to make a congenic inbred strain, homozygous for both gene mutations. Animals were housed under standard white cyclic lighting and had free access to food and water. Animal protocols compliant with §4 of the German law of animal protection were reviewed and approved by the Tübingen University committee on animal protection (Einrichtung für Tierschutz, Tierärztlicher Dienst und Labortierkunde, Registration No. AK02/19M, AK01/20M AK05/22M).

### Analysis of RNA-Seq, scRNA-seq data and differential expression

The mRNA expression comparison between *rd1* and *wt* mouse employed datasets downloaded from the GSE62020 database [27], while single-cell RNA expression analysis used data from the GSE212183 database [28]. To characterize *rd1*, we performed a differential analysis (fold change_J>_J1.2, *p* <_J0.05) comparing *rd1* to *wt* by using R language, the “limma” package. Heatmap and volcano plot were used to visualize Ca^2+^-related genes using the packages “pheatmap” and “ggplot2”. To infer functional annotations of *rd1* genes, gene ontology (GO) [m5.go.v2022.1.Mm.symbols.gmt] of differentially expressed genes (DEGs) was supplemented by gene set enrichment analysis (GSEA; version 4.2.2). The statistical significance was defined as false discovery rate (FDR) < 0.05, and the overrepresentation of indicated gene ontology (GO) gene sets in the ranked gene lists were presented by the normalized enrichment score (NES). GO enrichment analyses were conducted for the selected common DEGs using “GOplot” and “enrichplot” packages, *p* < 0.05. Additionally, the software packages “ggpubr”, “corrplot”, “fmsb” and “ggalluvial” were used for box plot, balloon plot, deviation plot, correlation plot, radar plot, and alluvial diagram, respectively.

### Retinal explant culture

To study the effects of various drugs on photoreceptor enzyme activities and cell death, *wt*, *rd1*, and *rd1*Cngb1^−/−^* retinas were explanted at postnatal day 5 (P5). The explants were cultured on a polycarbonate membrane (83.3930.040; 0.4 µm TC-inserts, SARSTEDT, Hildesheim, Germany) with complete medium (Gibco R16 medium with supplements) [29]. After explantation, the complete R16 medium was changed every two days with treatment. The two retinas obtained from a single animal were split across different experimental groups so as to maximize the number of independent observations acquired per animal. Cultures were treated with 10 µM BAPTA-AM (ab120503; Abcam, Cambridge, UK), 50µM L-cis-diltiazem (ab120532; Abcam), 20 µM CM4620 (HY-101942; MedChemExpress, Sollentuna, Sweden), 40 µM SN-6 (HY-107658; MedChemExpress), 100 µM D-cis-diltiazem (ab120260; Abcam), 10 µM TTA-A2 (HY-111828; MedChemExpress), 15 µM DS5565 (HY-108006; MedChemExpress), and 1 µM NA-184 (kindly provided by Michel Baudry, Western University, Pomona, CA, USA), respectively. In these treatments, all compounds were dissolved in DMSO at a final medium concentration of no more than 0.1% DMSO. Cultures were ended at P11 (*rd1* short-term cultures), P23 (*rd1* long-term treatment) and P17 (*rd1*Cngb1^-/-^*) by either fixation with 4% paraformaldehyde (PFA) or without fixation and direct freezing in liquid N_2_. Explants were embedded in Tissue-Tek (Sakura Finetek Europe B.V., Alphen aan den Rijn, The Netherlands) and sectioned (14 µm) in a cryostat (Thermo Fisher Scientific, CryoStar NX50 OVP, Runcorn, UK).

### TUNEL staining

TUNEL (terminal deoxynucleotidyl transferase dUTP nick end labelling) assay kit (Roche Diagnostics, Mannheim, Germany) was used to label dying cells. Histological sections from retinal explants were dried and stored at −20 °C. The sections were rehydrated with phosphate-buffered saline (PBS; 0.1 M) and incubated with proteinase K (1.5 µg/µL) diluted in 50 mM TRIS-buffered saline (TBS; 1 µL enzyme in 1 mL TBS) for 15 min. This was followed by 3 times 5 min TBS washing and incubation with blocking solution (10% normal goat serum, 1% bovine serum albumin, and 1% fish gelatine in phosphate-buffered saline with 0.03% Tween-20). TUNEL staining solution was prepared using 21 parts of blocking solution, 18 parts of TUNEL labelling solution, and 1 part of TUNEL enzyme. After blocking, the sections were incubated with TUNEL staining solution overnight at 4 °C. Finally, sections were washed 2 times with PBS, mounted using mounting medium with DAPI (ab104139; Abcam), and imaged by microscopy.

### Calpain activity assay

This assay allows resolving the overall calpain activity *in situ* on unfixed tissue sections. Retinal tissue sections were incubated and rehydrated for 15 min in a calpain reaction buffer (CRB) (5.96 g HEPES, 4.85 g KCl, 0.47 g MgCl_2_, and 0.22 g CaCl_2_ in 100 mL ddH_2_O; pH 7.2) with 2 mM dithiothreitol (DTT). Tissue sections were incubated for 3 h at 37 °C in CRB with tBOC-Leu-Met-CMAC (25 µM; A6520; Thermo Fisher Scientific, OR, USA). Then, sections were washed with PBS and incubated with ToPro (1:1000 in PBS, Thermo Fisher Scientific) for 15 min. Afterwards, tissue sections were washed twice in PBS (5 min) and mounted using Vectashield without DAPI (Vector Laboratories Inc., Burlingame, CA, USA) for immediate visualization by microscopy.

### PARP activity and PAR staining

The PARP *in situ* activity assay is based on the incorporation of a fluorescent NAD^+^ analogue and allows resolving the overall PARP enzyme activity on unfixed tissue sections [30]. Such sections were incubated and rehydrated for 10 min in PBS. The reaction mixture (10 mM MgCl_2_, 1mM dithiothreitol, and 50 μM 6-Fluo-10-NAD^+^ (Cat. Nr.: N 023; Biolog, Bremen Germany) in 100 mM Tris buffer with 0.2% Triton X100, pH 8.0) was applied to the sections for 3 h at 37 °C. After three 5 min washes in PBS, sections were mounted in Vectashield with DAPI (Vector Laboratories) for subsequent microscopy.

For the detection of PAR, we used an immunostaining enhanced with 3,3′-diaminobenzidine (DAB) staining. The procedure is initiated by quenching of endogenous peroxidase activity using 40% MeOH and 10% H_2_O_2_ in PBS with 0.3% Triton X-100 (PBST) in retinal tissue sections for 20 min. Sections were further incubated with 10% normal goat serum (NGS) in PBST for 30 min, followed by anti-PAR antibody (1:200; ALX-804-220-R100; Enzo Life Sciences, Farmingdale, NY, USA) incubation overnight at 4 °C. Incubation with the biotinylated secondary antibody (1:150, Vector in 5% NGS in PBST) for 1 h was followed by the Vector ABC Kit (Vector Laboratories, solution A and solution B in PBS, 1:150 each) for 1 h. DAB staining solution (0.05 mg/mL NH_4_Cl, 200 mg/mL glucose, 0.8 mg/mL nickel ammonium sulphate, 1 mg/mL DAB, and 0.1 vol. % glucose oxidase in phosphate buffer) was applied evenly, incubated for precisely 3 min, and immediately rinsed with phosphate buffer to stop the reaction. Sections were mounted in Aquatex (Merck, Darmstadt, Germany).

### Sirtuin/HDAC activity assay

This assay allows detecting overall HDAC activity *in situ* on fixed tissue sections and is based on an adaptation of the FLUOR DE LYS®-Green System (Biomol, Hamburg, Germany). Retinal sections were exposed to 50 µM FLUOR DE LYS®-SIRT1 deacetylase substrate (BML-Kl177-0005; ENZO, New York, USA) with 2 mM NAD^+^ (BML-KI282-0500; ENZO) in assay buffer (50 mM Tris/HCl, 137 mM NaCl; 2.7 mM KCl; 1mM MgCl2; pH 8.0) for 3 h at 37 °C. Sections were then washed in PBS and fixed in methanol at – 20 °C for 20 min. Slides were mounted with FLUOR DE LYS® developer II concentrate (BML-KI176-1250; Enzo, New York, USA) diluted 1:5 in assay buffer overnight for subsequent microscopy.

### Immunohistochemistry for calpain-1/2 and NCXs

Sections were rehydrated with PBS for 15 min and then incubated with a blocking solution (10% NGS, 1% BSA, and 0.3% PBST) for 1 h. The primary antibodies, rabbit-anti-calpain-2 (1:200; ab39165; Abcam), rabbit-anti-calpain-1 (1:100; ab39170; Abcam), rabbit-anti-NCX1 (1:200; LS-B15461; LifeSpan BioScience, Washington, USA), rabbit-anti-NCX2 (1:200; BS-1997R; BIOSS; Massachusetts, USA) and mouse-anti-NCX3 (1:100; NB120-2869; Novus Biologicals, Colorado, USA) were diluted in blocking solution and incubated overnight at 4 °C rinsing with PBS for 3 times 10 min each; this was followed by incubation with the secondary antibodies, goat-anti-rabbit AlexaFluor488 (1:400; A11034; Molecular Probes; Oregon, USA), goat-anti-rabbit AlexaFluor568 (1:300; A11036; Molecular Probes), and goat-anti-mouse AlexaFluor568 (1:500; A11031; Molecular Probes), for 1 h. The sections were further rinsed with PBS for 3 times 10 min each and mounted with mounting medium with DAPI (Abcam).

### Microscopy and image analysis in retinal cultures

Images of organotypic explant cultures were captured using a Zeiss Imager Z.2 fluorescence microscope, equipped with ApoTome 2, an Axiocam 506 mono camera, and HXP-120V fluorescent lamp (Carl Zeiss Microscopy, Oberkochen, Germany). Excitation (λExc.)/emission (λEm.) characteristics of filter sets used for different fluorophores were as follows (in nm): DAPI (λExc. = 369 nm, λEm = 465 nm), AF488 (λExc. = 490 nm, λEm = 525 nm), AF568 (λExc. = 578 nm, λEm = 602 nm), and ToPro (λExc. = 642 nm, λEm = 661 nm). The Zen 2.3 blue edition software (Zeiss) captured images (tiled and z-stack, 20× magnification). Sections of 14 µm thickness were analysed using 12–15 Apotome Z-planes. For quantification of positive cells in ONL, we proceeded as follows: The number of cells in six different rectangular ONL areas was counted manually based on the number of DAPI-stained nuclei, and used to calculate an average ONL cell size. This average ONL cell size was used to calculate the total number of cells in a given ONL area. The percentage of positive cells was then calculated by dividing the absolute number of positive cells by the total number of ONL cells. ONL thickness was determined by manual counts on DAPI stained retinal sections. For each count, a vertical column was placed on nine different positions of the section and the end-to-end distances of the photoreceptor layer in each of these positions were recorded manually. The nine individual counts were averaged to give the mean value of photoreceptor thickness for one explant.

### Statistical analysis

Two-way comparisons were analysed using Student’s *t*-test. Multiple comparisons were made using a one-way analysis of variance (ANOVA) test with Tukey multiple comparison test. Calculations were performed with GraphPad Prism 8 (GraphPad Software, La Jolla, CA, USA); *p* < 0.05 was considered significant; Levels of significance were as follows: *, *p* < 0.05; **, *p* < 0.01; ***, *p* < 0.001; ****, *p* < 0.0001. Data in figure 8A was normalized by linear scaling according to the formula: χ*_scaled_=*χ*-*χ*_min_/*χ*_max_-*χ*_min,_* using SPSS Statistics 26 (IBM, Armonk, New York, USA). The figures were prepared using Photoshop 2022 and Illustrator 2022 (Adobe, San Jose, CA, USA). Bioinformatic analyses were performed by R software (Version 4.0.1). The diagram was created with BioRender.com.

## RESULTS

### Ca^2+^-related genes and pathways during the retinal degeneration of *rd1*

To investigate whether and to what extent Ca^2+^ was involved in the progression of *rd1* retinal cell death, we performed RNA-seq to assess gene expression differences between *rd1* and *wt* mice. The GSE62020 database (accessed June 2023) was analysed to identify differentially expressed genes (DEGs). A fold change > 1.2 and a *p*-value < 0.05 were considered to indicate significant changes. Compared to *wt*, *rd1* retina expressed 667 up-regulated genes and – coincidentally – 667 down-regulated genes at P13. Among these, 89 DEGs were found to be associated with Ca^2+^-signalling, of which 32 were down-regulated and 57 were up-regulated in *rd1* retina (Figure 1A). Gene set enrichment analysis (GSEA) revealed four gene ontology (GO) terms involved in Ca^2+^-signalling (Figure 1B, FDR < 0.05). Ten Ca^2+^-related biology processes (BPs) and molecular functions (MFs) were identified by DEGs GO enrichment analysis (Figure 1C, *p* < 0.05).

**Figure 1.**
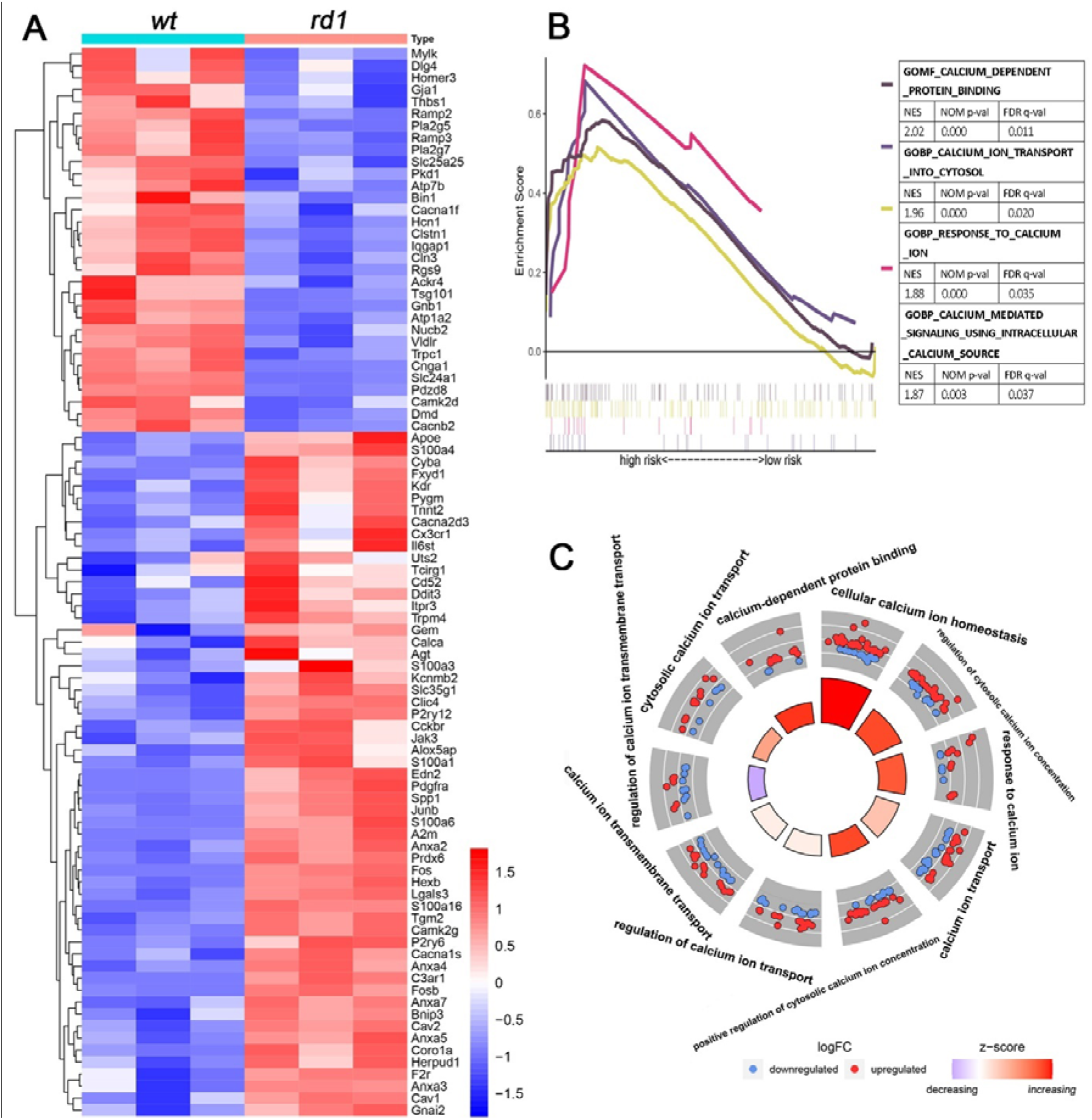
Whole tissue RNA-seq analysis highlights Ca^2+^-signalling-related changes in *rd1* retina. **A**) Heatmap for post-natal day 13 RNA-Seq data comparing the expression of Ca^2+^-related differentially expressed genes (DEGs) between *rd1* and wild-type (*wt*) retina. In this group of genes, 32 were down– and 57 were up-regulated in *rd1* retina. **B**) Gene set enrichment analysis (GSEA) showing upregulation of four Ca^2+^-related gene ontology (GO) pathways in *rd1* retina (false discovery rate, FDR < 0.05). **C**) Circle plot showing 10 different Ca^2+^-related GO terms (including biological processes (BPs) and molecular functions (MFs)) enriched in DEGs in the *rd1* situation (*p* < 0.05).

### Ca^2+^ plays an important role during photoreceptor degeneration

To further investigate Ca^2+^-related pathways specifically in degenerating *rd1* rod photoreceptors, we performed single-cell (sc) RNA-seq analysis to assess transcriptional differences between *rd1* and *wt* mice. A visualization in the form of a volcano plot showed the genes up– or down-regulated in rod photoreceptors from GSE212183. In total, 610 DEGs were found to be differentially expressed between *rd1* and *wt* rod photoreceptors at P13, where 336 DEGs were up-regulated and 274 were down-regulated (Figure 2A). Some of the DEGs were linked to Ca^2+^-related biological processes (BPs), according to GO enrichment analysis (Figure 2B). There were 780 DEGs found in *rd1* cone photoreceptors at P13 (Figure S1A). However, none of the cone DEGs were enriched in Ca^2+^-related GO terms.

**Figure 2.**
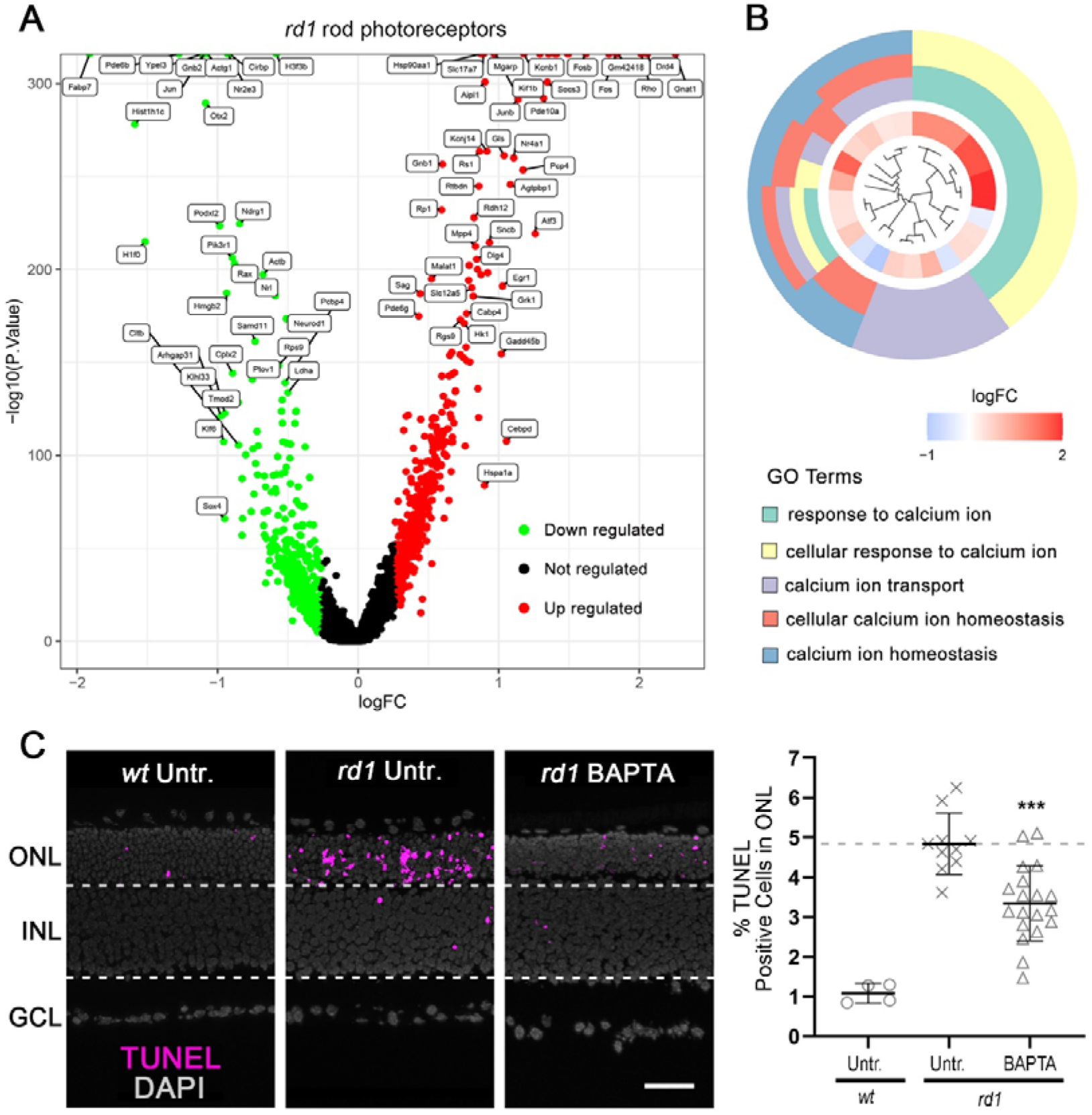
Rod photoreceptor scRNA-Seq and Ca^2+^ chelation reveals critical role for Ca^2+^-signalling in *rd1* cell death. **A**) Volcano plot for scRNA-Seq data showing differentially expressed genes (DEGs) in *rd1* and wild-type (*wt*) rod photoreceptors. **B**) Circle plot showing Ca^2+^-related GO terms enriched for DEGs in *rd1* rod photoreceptors (*p* < 0.05). **C**) The TUNEL assay labelled dying cells (magenta) in *wt* and *rd1* retinal explant cultures. DAPI (grey) was used as nuclear counterstain. Untreated (Untr.) *wt* and *rd1* retina were compared to retina treated with 10 µM BAPTA-AM (BAPTA). Scatter plot displaying percentage of TUNEL-positive cells in the outer nuclear layer (ONL). Statistical testing: Student’s *t*-test performed between *rd1* Untr. and 10 µM BAPTA-AM (BAPTA). Untr. *wt*: n = 4; Untr. *rd1*: 10; BAPTA *rd1*: 19; error bars represent SD; *** = *p* < 0.001. INL = inner nuclear layer, GCL = ganglion cell layer; scale bar = 50 µm.

In the following, we used the Ca^2+^ chelator BAPTA-AM in *rd1* organotypic retinal explants, with the aim of confirming the link between Ca^2+^ and photoreceptor degeneration. The TUNEL assay was used to quantify the numbers of dying cells in the outer nuclear layer (ONL). A dose-response for BAPTA-AM treatment on *rd1* explants was established and revealed 10 µM as a suitable concentration for further experiments (Figure S1B). In *wt* retinal explants, a relatively low number of ONL cells (1.08 % ± 0.21, n = 4) were positive for the TUNEL assay, when compared with their *rd1* counterparts (4.84 % ± 0.73, n = 10). BAPTA-AM treatment significantly reduced *rd1* ONL cell death to 3.34 % (± 0.92, n = 19, *p* < 0.001; Figure 2C). While BAPTA-AM treatment appeared to be well tolerated in *wt* retina (Figure S1E), it could not preserve photoreceptor viability in long-term treatment of *rd1* retina lasting until P23 (Figure S1C). In contrast, BAPTA-AM accelerated photoreceptor degeneration in explant cultures derived from *rd1*Cngb1^-/-^* double-mutant mice (Figure S1D), indicating that depletion of intracellular Ca^2+^ in photoreceptors lacking functional CNGC was also detrimental.

### Expression of Ca^2+^ permeable channels in retina and rod photoreceptors

A whole range of Ca^2+^-permeable channels could potentially contribute to increased intracellular Ca^2+^-levels in photoreceptors. Notably, the second messenger cGMP activates photoreceptor CNGC, leading to Na^+^ and Ca^2+^ influx and, indirectly, via ensuing changes in membrane polarization, to additional Ca^2+^-influx through VGCC [5]. High intracellular Na^+^ levels produced by CNGC could potentially reverse the directionality of NCX [31], resulting in Ca^2+^-influx instead of efflux. Moreover, high Ca^2+^, via the activation of calmodulin-dependent protein kinase 2 (CaMK2), could potentiate Ca^2+^-influx mediated by CRAC [32].

To assess the expression of these four groups of Ca^2+^-permeable channels, we used RNA-seq and scRNA–seq to screen for any possible expression changes connected to *rd1* photoreceptor degeneration. While in whole retina RNA-seq numerous changes were found in genes encoding for CNGC, VGCC, CRAC, and NCX isoforms, the most prominent expression changes appeared after P18, *i.e.* at a time when most *rd1* rod photoreceptors have already been lost (Figure 3A). At P13, RNA-seq data indicated that genes coding for VGCC were up-regulated while genes coding for CNGC were down-regulated (Figure 3B). In the next step, we therefore used scRNA-seq to assess gene expression changes in Ca^2+^-permeable channels specifically in *rd1* rod photoreceptors, in the P11 to P17 time-frame (Figure 3C; *cf*. Figure S2A for a corresponding scRNA-seq analysis for cone photoreceptors). At the peak of photoreceptor degeneration, at P13, we found an upregulation of genes coding for CNGC, VGCC, and NCX (Figure 3D), while CRAC genes appeared to be down-regulated. However, we also found an upregulation in the expression of the *Camk2g* gene, and CaMK2 may increase CRAC activity independent of CRAC gene expression changes (Figure S2B).

**Figure 3.**
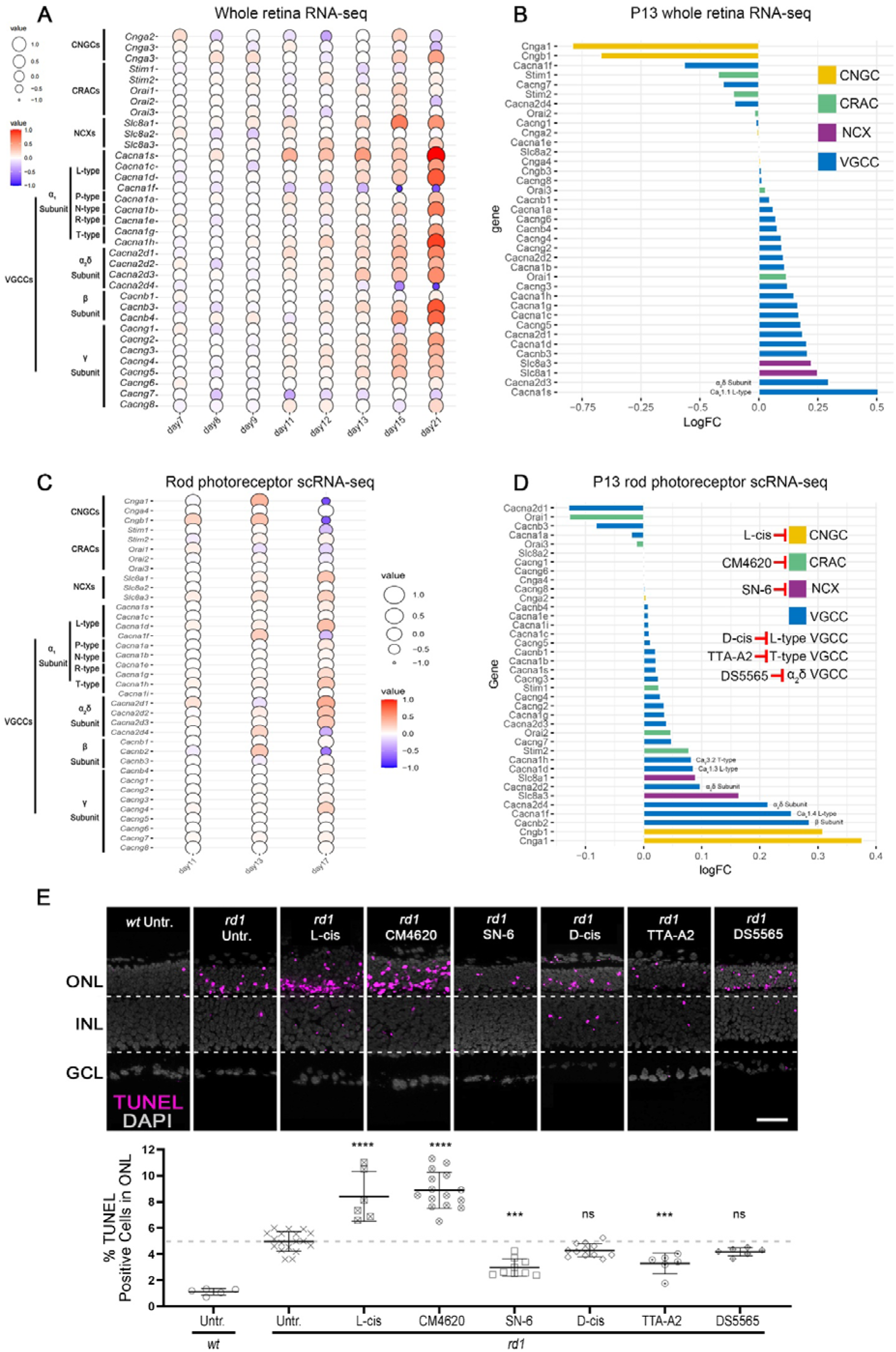
Ca^2+^-channels are differentially expressed in *rd1* retina and their inhibition can promote or reduce photoreceptor cell death. **A**) Balloon plot showing time-dependent whole-retina expression changes (post-natal day (P) 7 to P21) of cyclic nucleotide–gated channel (CNGC), Ca^2+^-release activated channel (CRAC), Na^+^/Ca^2+^ exchanger (NCX), and voltage-gated Ca^2+^ channel (VGCC). **B**) Deviation plot highlighting expression changes for CNGC, CRAC, NCX, and VGCC in P13 *rd1* whole retina. **C**) Balloon plot showing scRNA-seq data and time-dependent expression changes (post-natal day (P) 11 to P17) of CNGC, CRAC, NCX, and VGCC in *rd1* rod photoreceptors. **D**) Deviation plot highlighting expression changes for CNGC, CRAC, NCX, and VGCC in P13 *rd1* rod photoreceptors. **E**) TUNEL assay labelling dying cells (magenta) in *rd1* and wild-type (*wt*) retinal explant cultures. DAPI (grey) was used as a nuclear counterstain. Untreated (Untr.) *wt* and *rd1* retina were compared to retina treated with 50 µM CNGC inhibitor (L-cis diltiazem), 20 µM CRAC inhibitor (CM4620), 40 µM NCX inhibitor (SN-6), 100 µM L-type VGCC inhibitor (D-cis diltiazem), 10 µM T-type VGCC inhibitor (TTA-A2), and 15 µM α_2_δ subunit VGCC ligand (DS5565). Scatter plot showing percent TUNEL-positive cells in the outer nuclear layer (ONL). The dashed line indicates the *rd1* untr. situation, data points below this threshold indicate protective effects, data points above suggest destructive effects. Statistical testing: one-way ANOVA and Tukey’s multiple comparison post hoc test. Untr. *wt*: n = 5; Untr. *rd1*: 18; L-cis *rd1*: 6; CM4620 *rd1*: 15; SN-6 *rd1*: 9; D-cis *rd1*: 12; TTA-A2 *rd1*: 6; DS5565 *rd1*: 6; error bars represent SD; significance levels: *** = *p* < 0.001; **** = *p* < 0.0001. INL = inner nuclear layer, GCL = ganglion cell layer; scale bar = 50 µm.

Overall, the RNA-seq and scRNA-seq data suggested important changes in the expression of Ca^2+^– permeable channel-related genes in *rd1* photoreceptor degeneration. Still, from the RNA expression data alone it was not possible to deduce which type of Ca^2+^-permeable channel was causally involved in *rd1* cell death.

### Inhibition of Ca^2+^ permeable channels impact rod photoreceptor viability

To characterize the role of different Ca^2+^-permeable channels functionally, we employed an array of different channel inhibitors, including the CNGC inhibitor L-cis-diltiazem (L-cis), the CRAC inhibitor CM4620, the NCX inhibitor SN-6, the L-type VGCC inhibitor D-cis-diltiazem (D-cis), the T-type VGCC inhibitor TTA-A2, and the α_2_δ subunit VGCC ligand DS5565. Retinal explant cultures derived from *rd1* and *wt* animals were treated with these inhibitors and the effects analysed using the TUNEL assay to detect dying cells in the ONL. To assess a possible cross-reactivity of the inhibitors used, we also employed explant cultures derived from *rd1*Cngb1^-/-^*double-mutant mice, *i.e.* retinas in which rods lack functional CNGC. For L-cis and D-cis suitable drug concentrations for retinal treatments had been established previously [33]; for inhibitors where such data were missing (*i.e*. SN-6, TTA-A2, DS5565, CM4620), dose-response curves were generated to select appropriate treatment concentrations (Figures S2, S3).

As shown previously [14, 34], in *wt* retina the number of TUNEL positive cells in the ONL was significantly lower than in its *rd1* counterpart (Figure 3E, Table S1). Remarkably, treatment with both L-cis and CM4620 significantly increased the numbers of TUNEL positive cells in both *rd1* and *wt* ONL (Figure 3E, Figure S1E, Table S1). Conversely, SN-6 and TTA-A2 significantly decreased photoreceptor cell death in *rd1* retina (Figure 3E, Table S1). Also, in the retina of *rd1*Cngb1^-/-^* double-mutant mice SN-6 significantly decreased TUNEL positive cells (Figure S1D), yet, in *wt* retina ONL cell death was significantly increased by SN-6 (Figure S1E). Similar to BAPTA-AM, SN-6 did not show long-term protection in *rd1* ONL (Figure S1C). To assess the expression of NCX at the protein level, immunostaining was performed using antibodies directed against NCX1, NCX2, and NCX3. When compared to negative control, only NCX1 was found to be expressed in *rd1* and *wt* retina, notably in photoreceptor segments and Müller glial cells (Figure S3A).

The compounds D-cis and DS5565 targeting L-type and α_2_δ VGCC did not lead to significant reduction of TUNEL positive cells in *rd1* ONL (Figure 3E, Table S1).

Taken together, the experiments with Ca^2+^-permeable channel blockers confirmed that Ca^2+^-signalling was indeed important for photoreceptor viability. Contrary to expectations, activity of CNGC and CRAC appeared to have a pro-survival role, while activity of T-type VGCC and NCX was detrimental for *rd1* rod photoreceptors. The role of NCX appeared ambiguous, as its inhibition improved survival of *rd1* photoreceptors but promoted death in *wt* ones, indicating that NCX directionality might have been reversed in the *rd1* rod photoreceptors. Finally, L-type VGCC activity appeared to be unrelated to *rd1* degeneration.

### Calpain-2 contributes to *rd1* photoreceptor degeneration

The reduction of TUNEL-positive cells after Ca^2+^ chelation and Ca^2+^ permeable channel inhibition strongly suggested a link between high intracellular Ca^2+^-levels and *rd1* photoreceptor cell death. To further investigate Ca^2+^-induced cell death, we performed GSEA analysis on *rd1* P13 RNA-seq data, in which DEGs were enriched in the GO biological process (BP) pathways, including “Positive regulation of proteolysis” (GO: 0045862; normalized enrichment score (NES) = 1.78, *p* < 0.0001; Figure 4A), “Membrane protein proteolysis” (GO: 0033619; NES = 1.62, *p* = 0.003; Figure 4B), “Membrane protein ectodomain proteolysis” (GO: 0006509; NES = 1.6, *p* = 0.027; Figure 4C), and “Positive regulation of proteolysis involved in protein catabolic process” (GO: 1903052; NES = 1.6, *p* < 0.0001; Figure 4D). Previous studies had connected the activity of Ca^2+^-dependent calpain-type proteases to *rd1* retinal degeneration [35, 36] and we hypothesized that the neurodegenerative calpain-2 isoform [37] might be responsible for retinal cell death. Thus, we used the recently developed and highly specific calpain-2 inhibitor NA-184 [38] to treat *rd1* explant cultures. A dose-response curve for the effects of NA-184 was compiled (Figure S3D) and a concentration of 1 µM was chosen for further experiments. As before, in *wt* retina the numbers of TUNEL positive cells in the ONL were low (1.08 % ± 0.44, n = 4) when compared with untreated *rd1* retina (4.8 % ± 0.47, n = 7; Figure 4E). NA-184 treatment significantly reduced *rd1* ONL TUNEL positive cells (3.55 % ± 0.53, n = 7, *p* < 0.01; Figure 4E), implying a causative involvement of calpain-2 in the degenerative process.

**Figure 4.**
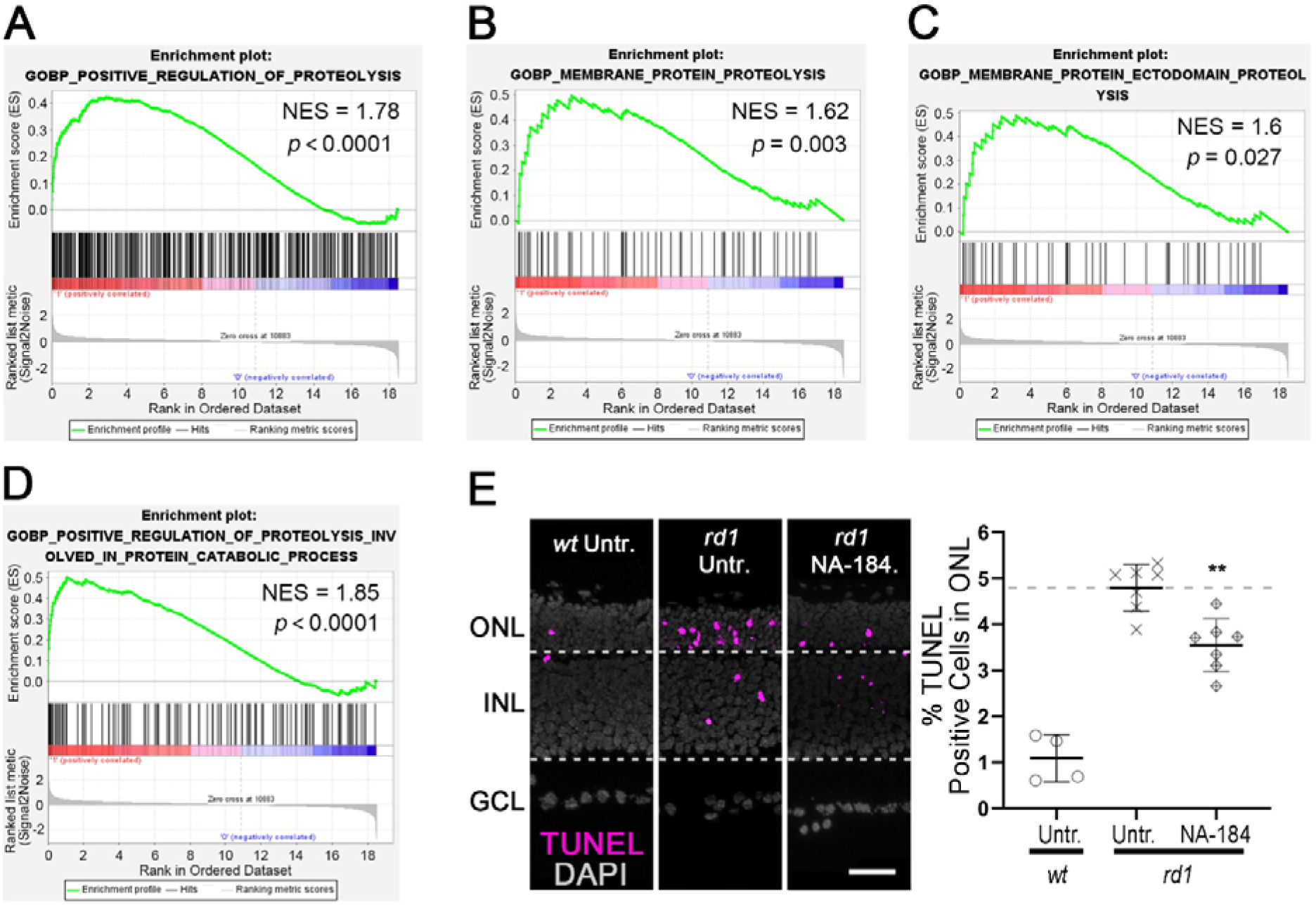
Ca^2+^-dependent proteolysis and calpain-2 are associated with *rd1* photoreceptor cell death. **A**) Gene set enrichment analysis (GSEA) showing enrichment of differentially expressed genes (DEGs) in the biological process (BP) “Positive regulation of proteolysis” (GO: 0045862). **B**) DEGs enriched in the BP “Membrane protein proteolysis” (GO: 0033619). **C**) DEGs enriched in the BP “Membrane protein ectodomain proteolysis” (GO: 0006509). **D**) DEGs enriched in the BP “Positive regulation of proteolysis involved in protein catabolic process” (GO:1903052). **E**) TUNEL assay labelled dying cells (magenta) in wild-type (*wt*) and *rd1* retinal explant cultures. DAPI (grey) was used as nuclear counterstain. Untreated (Untr.) *wt* and *rd1* retina compared to retina treated with the calpain-2 selective inhibitor NA-184. The scatter plot shows the percentage of TUNEL-positive cells in the outer nuclear layer (ONL). Statistical comparison: Student’s *t*– test performed between *rd1* Untr. and *rd1* NA-184. INL = inner nuclear layer, GCL = ganglion cell layer; scale bar = 50 µm.

### Calpain activity changes after interventions targeting intracellular Ca^2+^

To further study how Ca^2+^ chelation, activity of Ca^2+^ permeable channels, and calpain-2 inhibition regulated overall calpain, we investigated calpain activity using a general *in situ* activity assay and immunolabelling for activated calpain-1 and calpain-2. Calpain activity and calpain-2 activation were rather low in *wt* retina when compared with *rd1* retina (Figure 5A, C; Table S2A, C). BAPTA-AM significantly reduced both overall calpain activity and calpain-2 activation specifically (Figure 5A, C; Table S2A, C; dose-response curve shown in Figure S1B). However, L-cis and CM4620 significantly increased both calpain activity and calpain-2 activation (Figure 5A, C; Table S2A, C; dose response curve for CM4620 shown in Figure S2C). General calpain activity and calpain-2 activation were also significantly reduced after treatment with SN-6, D-cis, TTA-A2, and NA-184 (Figure 5A, C; Table S2A, C). Dose-response for SN-6, TTA-A2, and NA-184 on *rd1* calpain activity are shown in Figure S2D, Figure S3B, D, respectively. Interestingly, DS5565 significantly reduced overall calpain activity, while it did not decrease calpain-2 activation, when compared to *rd1* control (Figure 5A, C; Table S2A, C; dose response curve for DS5565 shown in Figure S3C).

**Figure 5.**
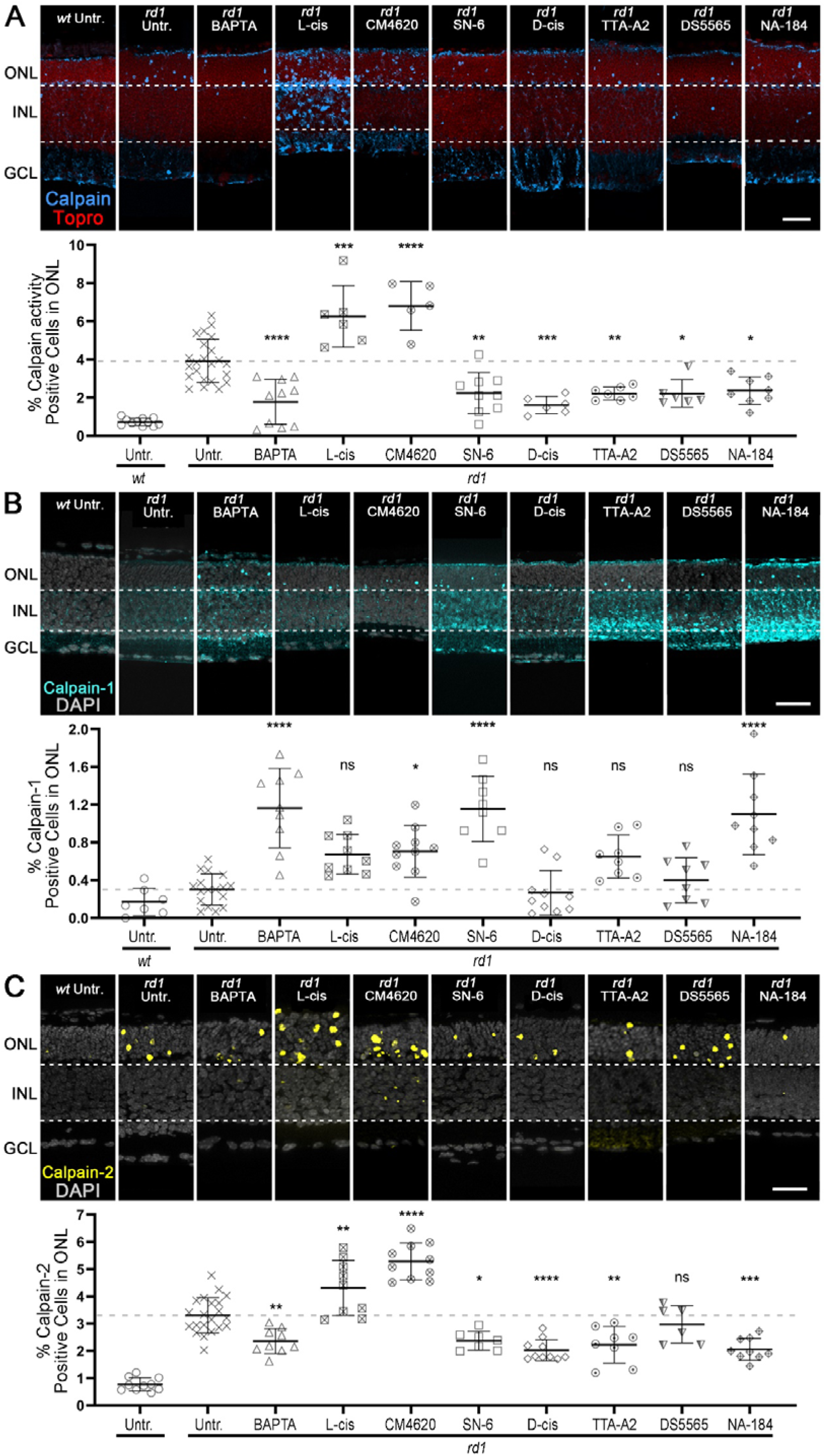
Ca^2+^-channel inhibitors differentially regulate general calpain activity, activation of calpain-1 and –2. **A**) Calpain activity assay (blue) was performed on wild-type (*wt*) and *rd1* retinal explant cultures. ToPro (red) was used as a nuclear counterstain. Untreated (Untr.) *wt* and *rd1* retina were compared to retina treated with BAPTA-AM, L-cis diltiazem, CM4620, SN-6, D-cis diltiazem, TTA-AS, DS5565, and NA-184. Scatter plot showing percent calpain activity positive cells in the outer nuclear layer (ONL). Untr. *wt*: n = 11; Untr. *rd1*: 23; BAPTA *rd1*: 10; L-cis *rd1*: 6; CM4620 *rd1*: 5; SN-6 *rd1*: 9; D-cis *rd1*: 6; TTA-A2 *rd1*: 7; DS5565 *rd1*: 6; NA-184 *rd1*: 8. **B**) Activated calpain-1 (cyan) immunostaining was performed in *wt* and *rd1* retinal explant cultures. DAPI (grey) was used as a nuclear counterstain. Untreated *wt* and *rd1* retina were compared to retina treated with compounds as in A. Scatter plot showing percent displaying calpain-1 activation. Untr. *wt*: n = 7; Untr. *rd1*: 17; BAPTA *rd1*: 9; L-cis *rd1*: 9; CM4620 *rd1*: 10; SN-6 *rd1*: 8; D-cis *rd1*: 10; TTA-A2 *rd1*: 8; DS5565 *rd1*: 8; NA-184 *rd1*: 9. **C**) Activated calpain-2 (yellow) immunostaining was performed in *rd1* and *wt* retinal explant cultures. DAPI (grey) was used as a nuclear counterstain. Untreated *wt* and *rd1* retina were compared to retina treated with compounds as in A. Scatter plot showing percent ONL cells displaying calpain-2 activation. Untr. *wt*: 10; Untr. *rd1*: 21; BAPTA *rd1*: 9; L-cis *rd1*: 9; CM4620 *rd1*: 10; SN-6 *rd1*: 7; D-cis *rd1*: 10; TTA-A2 *rd1*: 8; DS5565 *rd1*: 6; NA-184 *rd1*: 9;. Note the significant elevation of calpain activity/calpain-2 activation caused by the CNGC inhibitor L-cis diltiazem and the CRAC inhibitor CM4620. The opposite effect is observed for calpain-1 activation. Statistical testing: one-way ANOVA and Tukey’s multiple comparison post hoc test; significance levels: * = *p* < 0.05; ** = *p* < 0.01; *** = *p* < 0.001; **** = *p* < 0.0001; error bars represent SD; INL = inner nuclear layer, GCL = ganglion cell layer; scale bar = 50 µm.

In *wt* retina the numbers of calpain-1 activation in ONL were relatively similar when compared with that of *rd1* (*p* > 0.05; Figure 5B and Table S2B). BAPTA-AM significantly increased the percentage of activated calpain-1 in *rd1* ONL, while L-cis did not (Figure 5B; Table S2B). The number of activated calpain-1 positive cells in *rd1* ONL was significantly raised by CM4620 and SN-6, but not by D-cis, TTA-A2, and DS5565 (Figure 5B; Table S2B). Interestingly, the calpain-2 inhibitor NA-184 significantly increased calpain-1 activation in *rd1* retina (Figure 5B; Table S2B).

### Relationships between activation of calpain-1, calpain-2, and cell death

To further investigate the interactions between calpain-1, calpain-2, and cell death, we used the corresponding datasets to perform Spearman’s rank correlation coefficient analysis (Spearman analysis) by assessing the strength and direction of the monotonic relationship between two variables [39]. Here, calpain-1 activation in *rd1* ONL correlated negatively with calpain-2 activation (*R* = –0.62, *p* < 0.01, Figure 6A), and with TUNEL positive cells (*R* = –0.65, *p* < 0.0001, Figure 6B). However, the ratio of calpain-2/calpain-1 was positively correlated with TUNEL (*R* = 0.61, *p* < 0.01, Figure 6C). The correlation analysis thus suggested that calpain-1 activation was not connected to cell death, while calpain-2 activation was.

**Figure 6.**
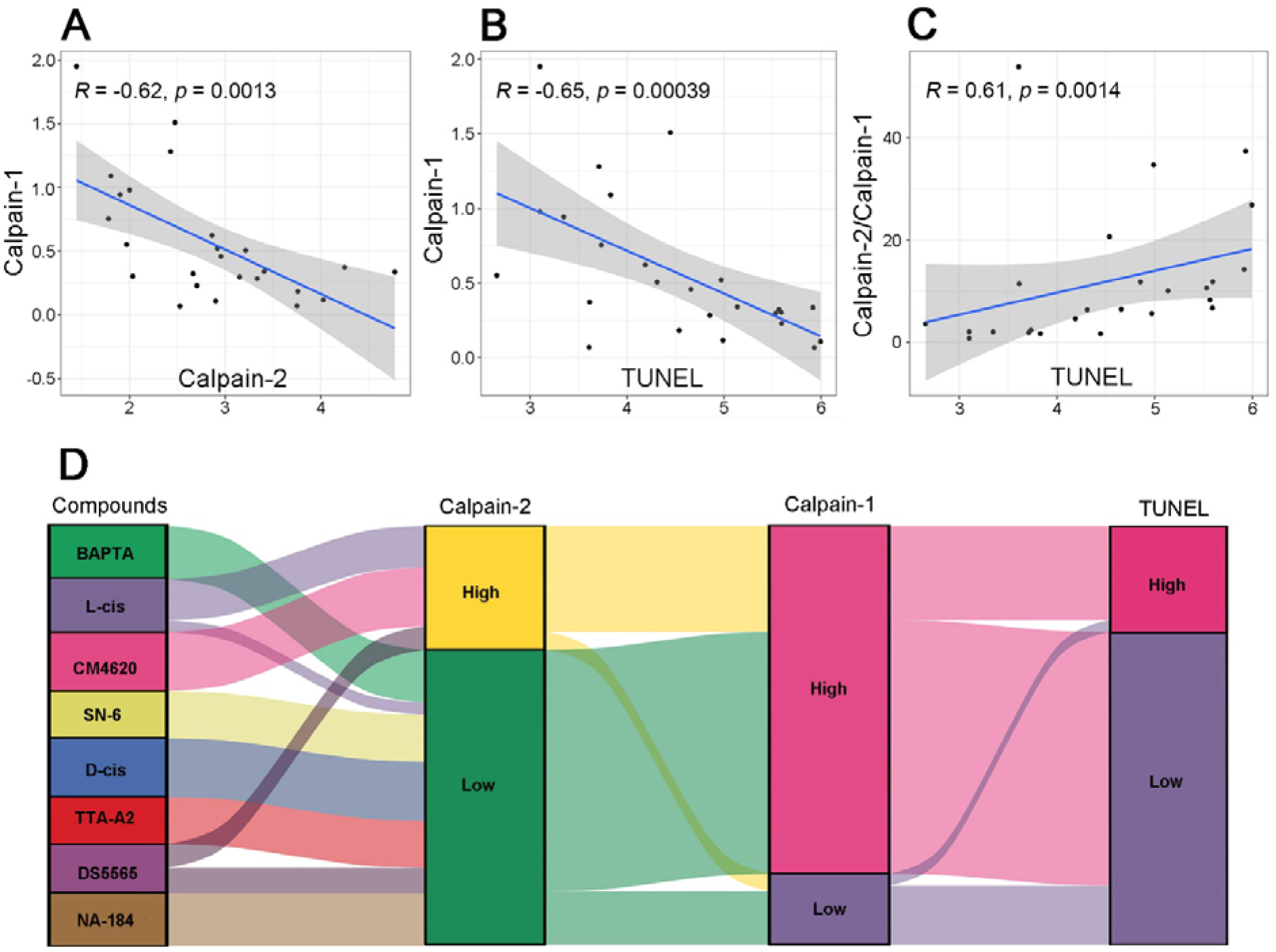
Differential correlation of calpain-1 and calpain-2 to photoreceptor cell death. **A**) Spearman analysis comparing the numbers of activated calpain-1 and calpain-2 in the outer nuclear layer (ONL) of untreated *rd1* retina and *rd1* retina treated with the calpain-2 inhibitor NA-184. **B**) Spearman analysis between the numbers of activated calpain-1 and dying, TUNEL positive cells in untreated or NA-184 treated *rd1* ONL. **C**) Spearman analysis between the ratio of calpain-2/calpain-1 and TUNEL positive cells in untreated or NA-184 treated *rd1* ONL. **D**) Alluvial diagram showing the relationship between activated calpain-2 and calpain-1, as well as TUNEL positive cells in *rd1* ONL after treatment with BAPTA-AM, L-cis, CM4620, SN-6, D-cis, TTA-AS, DS5565, and NA-184.

To assess general trends in the relationships between calpain-1, calpain-2, and cell death, we used an alluvial diagram where the numbers of activated calpain-1, –2, and TUNEL positive cells in the ONL of the *rd1* control group were used as baseline (Figure 6D). This representation indicated that in *rd1* retinal explant cultures treatment with BAPTA-AM, SN-6, D-cis, TTA-A2, DS5565, and NA-184 led to low calpain-2 activation in ONL when compared to untreated *rd1* (Figure 6D). In contrast, L-cis and CM4620 treatments were related to high calpain-2 activation. Most of the high calpain-2 activation was related to low calpain-1 activation which, in turn, was related to a large number of TUNEL positive cells in ONL (Figure 6D). Taken together, this analysis suggested that calpain-1 activation was unrelated to cell death while calpain-2 activation was closely connected to cell death.

### Effect of Ca^2+^ and calpain-2 on enzymatic activities of PARP and sirtuin

PARP was previously reported to be connected to photoreceptor cell death [4]. To dissect the relationship of Ca^2+^, calpain, and PARP, we performed an *in situ* PARP activity assay and immunostaining for poly(ADP-ribose) (PAR), *i.e*. the product of PARP activity. While PARP activity and PAR-positive cells were infrequent in *wt* retina when compared with *rd1* retina (Figure 7A, B; Table S3A, B), treatment with BAPTA-AM significantly reduced both PARP activity and PAR generation in *rd1* ONL (Figure 7A, B; Table S3A, B; BAPTA-AM dose-response in Figure S1B), indicating a Ca^2+^-dependent mechanism for PARP activation. However, the Ca^2+^-channel blockers L-cis and CM4620 significantly increased both PARP activity and PAR in *rd1* ONL (Figure 7A, B; Table S3A, B; CM4620 dose-response in Figure S2C). In contrast, PARP activity and PAR positive cells in *rd1* ONL were significantly reduced after treatment with SN-6, D-cis, TTA-A2, and DS5565 (Figure 7A, B; Table S3A, B; dose response curves for SN-6, TTA-A2, DS5565 shown in Figures S2D, S3B, C, respectively). The calpain-2 inhibitor NA-184 neither decreased PARP activity nor PAR generation in *rd1* ONL (Figure 7A, B; Table S3A, B; NA-184 dose-response in Figure S3D).

**Figure 7.**
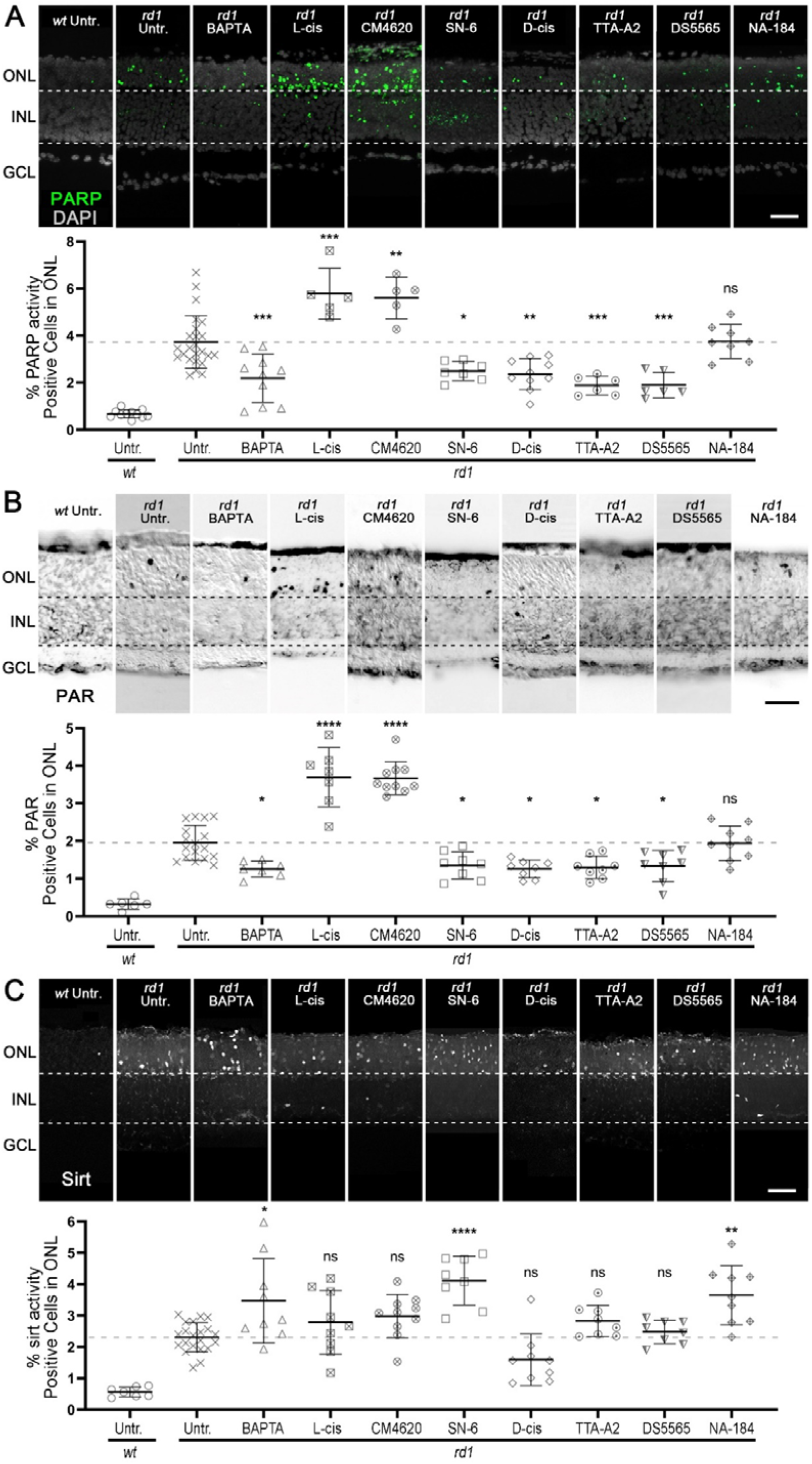
Ca^2+^-signalling affects PARP activity, PAR generation, and sirtuin activity. **A**) PARP activity assay (green) was performed in *rd1* and wild-type (*wt*) retinal explant cultures. DAPI (grey) was used as a nuclear counterstain. Untreated (Untr.) *rd1* and *wt* retina were compared to *rd1* retina treated with BAPTA-AM, L-cis, CM4620, SN-6, D-cis, TTA-AS, DS5565, and NA-184. The scatter plot shows percentage of PARP activity positive cells in the outer nuclear layer (ONL). Untr. *wt*: n = 11; Untr. *rd1*: 24; BAPTA *rd1*: 10; L-cis *rd1*: 5; CM4620 *rd1*: 5; SN-6 *rd1*: 7; D-cis *rd1*: 10; TTA-A2 *rd1*: 6; DS5565 *rd1*: 6; NA-184 *rd1*: 8. **B**) PAR staining (black) was performed in *rd1* and *wt* retinal explant cultures. Untreated *wt* and *rd1* retina were compared to drug-treated retina as in A. Untr. *wt*: n = 6; Untr. *rd1*: 17; BAPTA *rd1*: 7; L-cis *rd1*: 7; CM4620 *rd1*: 10; SN-6 *rd1*: 8; D-cis *rd1*: 8; TTA-A2 *rd1*: 8; DS5565 *rd1*: 8; NA-184 *rd1*: 9. **C**) Sirtuin activity assay (white) was performed in *rd1* and *wt* retinal explant cultures. Untreated *rd1* and *wt* retina were compared to drug-treated retina as in A. Untr. *wt*: n = 7; Untr. *rd1*: 21; BAPTA *rd1*: 9; L-cis *rd1*: 9; CM4620 *rd1*: 10; SN-6 *rd1*: 8; D-cis *rd1*: 9; TTA-A2 *rd1*: 8; DS5565 *rd1*: 8; NA-184 *rd1*: 9. Statistical testing: one-way ANOVA with Tukey’s multiple comparison post hoc test performed between *rd1* explant cultures. Error bars represent SD; ns = *p* > 0.05; * = *p* < 0.05; ** = *p* < 0.01; *** = *p* < 0.001; **** = *p* < 0.0001. INL = inner nuclear layer, GCL = ganglion cell layer. Scale bar = 50 µm.

The histone deacetylase (HDAC) sirtuin-1 (SIRT1) was suggested to be indirectly regulated by PARP-dependent consumption of NAD^+^ [40]. Thus, we investigated sirtuin activity by using a general HDAC *in situ* activity assay based on deacetylation of a SIRT1 specific substrate. In *rd1* retina, the number of HDAC or sirtuin activity positive cells in the ONL was higher compared to *wt* control (Figure 7C; Table S3C). BAPTA-AM further significantly increased sirtuin activity in *rd1* ONL (Figure 7C; Table S3C). Treatment with L-cis and CM4620 did not change the numbers of sirtuin activity positive cells, as compared to untreated *rd1*, while sirtuin activity was increased by SN-6 (Figure 7C; Table S3C). Sirtuin activity did not rise after treatment with D-cis, TTA-A2, and DS5565 (Figure 7C; Table S3C), but unexpectedly, NA-184 significantly increased the percentage of sirtuin positive cells in *rd1* ONL (Figure 7C; Table S3C).

### Enzyme activity patterns are altered by changes in Ca^2+^ or inhibition of calpain-2

**Figure 8.**
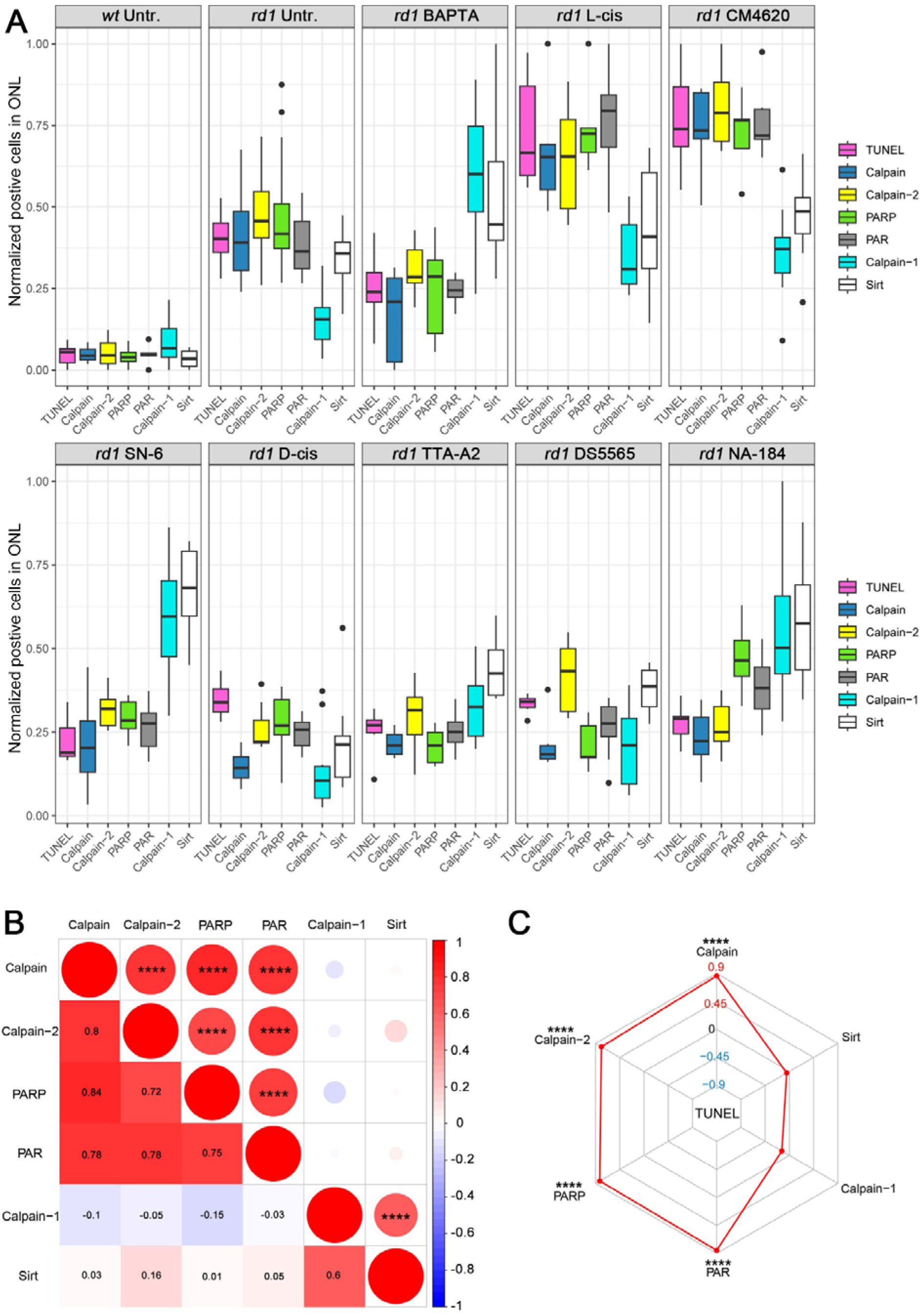
Enzymatic signatures for *rd1* photoreceptor degeneration and Spearman analysis. **A**) Comparison of enzymatic markers across different experimental treatments. Normalized cell numbers positive for TUNEL (magenta), calpain activity (calpain, blue), activated calpain-2 (yellow), PARP activity (PARP, green), PAR (black), activated calpain-1 (cyan), and sirtuin (Sirt) activity (white). **B**) Spearman analysis of enzymatic markers in photoreceptors (calpain activity, calpain-1, calpain-2, PARP activity, PAR, Sirtuin). The asterisks in circles show statistical significance, numbers in squares present the *R^2^*. **C**) Radar plot for Spearman analysis revealing the correlation between different enzymatic markers and the TUNEL assay. Note that while cell death (TUNEL) was strongly associated with general calpain activity, calpain-2 activation, PARP activity, and PAR, it was negatively correlated with calpain-1 activation and Sirtuin activity.

To compare the various enzymatic markers with each other, we normalized the experimental data by linear scaling, such that the lowest values were set to zero while the highest values were set to one. In *wt* retina, all enzyme activity markers were generally low when compared with the *rd1* untreated group (Figure 8A). In untreated *rd1*, the number of ONL cells showing calpain-1 activation was lower than calpain activity, calpain-2 activation, PARP activity, PAR positive cells, and sirtuin activity (Figure 8A). Photoreceptor degeneration was significantly reduced by treatments with BAPTA-AM, SN-6, TTA-A2, and NA-184, while calpain-1 activation and sirtuin activity were relatively high, compared to other markers (Figure 8A). In contrast, the treatments with L-cis and CM4620, which increased photoreceptor death, increased calpain activity, calpain-2 activation, PARP activity, and PAR-positive cells more than calpain-1 activation and sirtuin activity (Figure 8A). In the D-cis and DS5565 treated groups, which did not affect photoreceptor viability, calpain-1 activation in ONL was generally lower than calpain-2 activation (Figure 8A). Taken together, the activity patterns observed might constitute enzymatic signatures characteristic for either retinal degeneration or protection.

To go into a deeper analysis of these activity patterns triggered by alterations in Ca^2+^-influx, we performed a Spearman analysis (Figure 8B). This analysis excluded data from the NA-184 treatment since this calpain inhibitor would not *per se* change Ca^2+^-influx. General calpain activity and calpain-2 activation was positively correlated with PARP activity and PAR (*R* > 0.5, *p* < 0.0001). Calpain-1 activation and sirtuin activity were also positively corelated with each other (*R* > 0.5, *p* < 0.0001). A radar plot was used to show the relationship of cell death and enzymatic signatures after Spearman analysis (Figure 8C). This illustrated that TUNEL positive cells in *rd1* ONL were positively corelated with calpain activity, calpain-2 activation, PARP activity and PAR (*R* > 0.5, *p* < 0.0001, Figure 8C). However, TUNEL positive cells were anti-correlated with sirtuin activity and calpain-1 activation. Taken together, these analyses suggest that calpain-1 activation and sirtuin activity were associated with photoreceptor survival, as opposed to general calpain activity, calpain-2, PARP, and PAR, which were strongly connected to cell death.

## DISCUSSION

Ca^2+^ overload mediated by CNGC and VGCC has repeatedly been connected to cGMP-induced photoreceptor death in IRD models [13, 41, 42]. However, a number of studies using inhibitors of Ca^2+^-permeable channels have reported contradictory effects [43, 44], such that the precise role of Ca^2+^ in photoreceptor degeneration is still unclear. Our present work confirms that an imbalance in intracellular Ca^2+^ can cause photoreceptor degeneration and that blocking Ca^2+^-influx can be neuroprotective. Notably, our data suggests the Ca^2+^-permeable channels T-type VGCC, and possibly also NCX, as therapeutic targets for the treatment of IRD (Figure 9). Similarly, inhibition of the Ca^2+^-activated proteolytic enzyme calpain-2 was also found to be neuroprotective. As opposed to calpain-2, calpain-1 activity was linked to lower cell death rates. Overall, our study indicates that both too high and too low intracellular Ca^2+^ levels are detrimental to photoreceptor viability.

**Figure 9.**
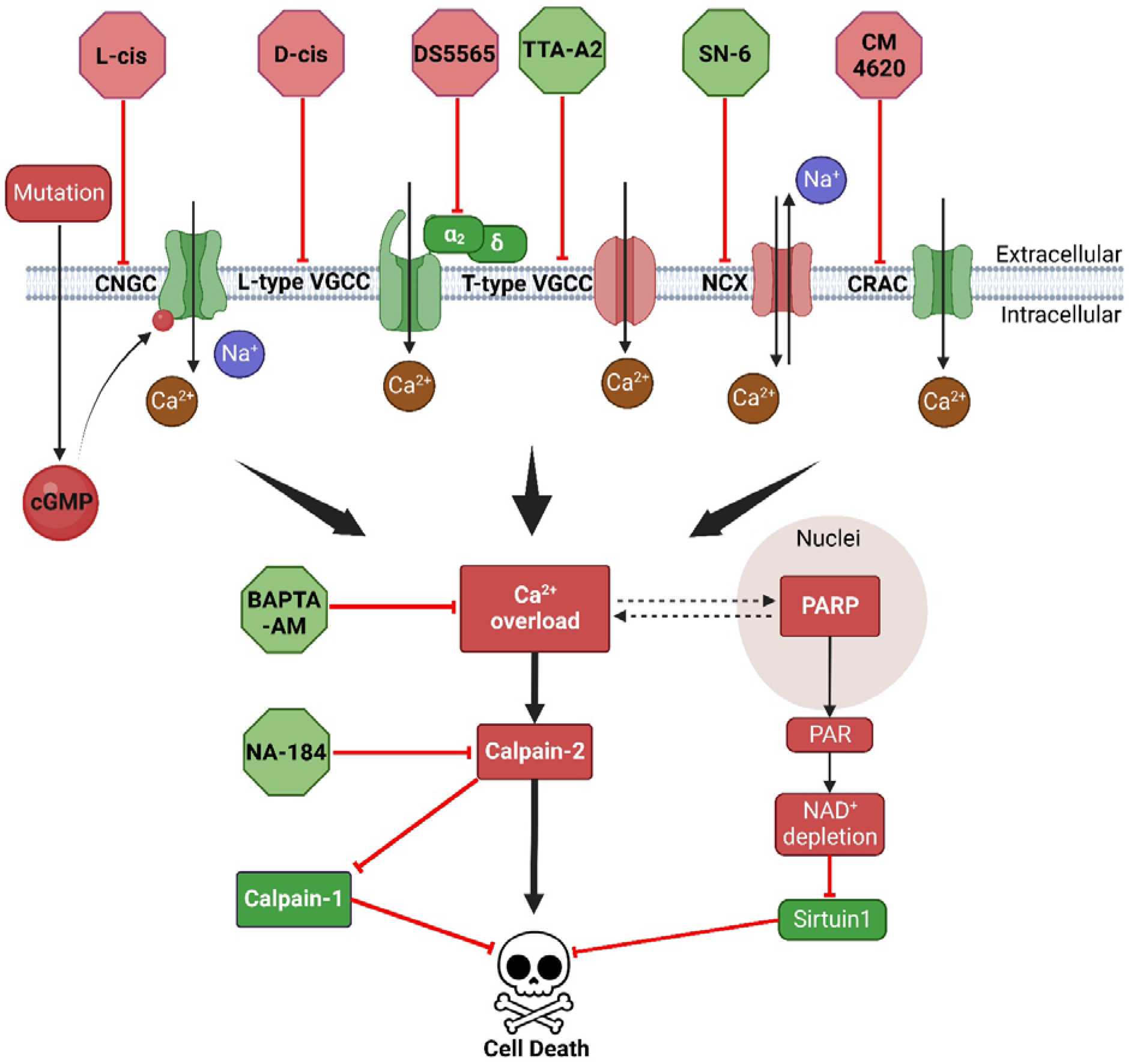
Experimental interventions and their relation to Ca^2+^ signalling in cGMP-dependent *rd1* degeneration. In *rd1* photoreceptors, the *Pde6b* mutation induces cGMP accumulation, which activates cyclic-nucleotide-gated channels (CNGC), leading to Na^+^ and Ca^2+^ influx. CNGC-dependent depolarization activates voltage-gated Ca^2+^ channel (VGCC) and may reverse directionality of Na^+^/Ca^2+^ exchanger (NCX), both leading to more Ca^2+^-influx. Additional Ca^2+^-influx could be mediated by Ca^2+^ release activated channel (CRAC). All these Ca^2+^ permeable channels contribute to intracellular Ca^2+^ overload, which, in turn, may activate calpain-2 directly and poly(ADP-ribose)-polymerase (PARP) indirectly. On the one hand, over-activated calpain-2 increases proteolysis of neuroprotective calpain-1. On the other hand, Ca^2+^-dependent activation of PARP may positively feedback on Ca^2+^-influx via NAD^+^ depletion, which may reduce the protective activity of NAD^+^-dependent sirtuins. Eventually, the activities of calpain-2 and PARP, triggered by high intracellular Ca^2+^-levels promote photoreceptor cell death. The drugs used in this study and their targets are indicated. Red colour indicates destructive processes and drugs, while green labelled proteins and compounds promote photoreceptor survival.

### Ca^2+^ homeostasis is critical for the survival of retinal photoreceptors

In different cell types, the second messenger Ca^2+^ is involved in the regulation of a diverse range of cellular processes, including fertilization, metabolism, transcription, and cell death [45]. This diversity of Ca^2+^-signalling related processes requires a precise control of intracellular Ca^2+^ levels, which in a healthy cell are maintained at below 100 nM *vs*. extracellular levels exceeding 2 mM [45]. In photoreceptors, the over-activation of CNGC caused by high intracellular cGMP levels leads to an influx of Na^+^ and Ca^2+^, depolarizing the cell and activating VGCC, causing further Ca^2+^ influx [5, 42]. However, whether Ca^2+^ is responsible for photoreceptor degeneration is debated, as some reports indicate that blocking Ca^2+^-permeable channels may be neuroprotective [16, 17, 46], while other studies propose the opposite [18, 19]. Our bioinformatic analysis of RNA-seq data obtained at the peak of *rd1* photoreceptor cell death (P13) revealed more than 80 DEGs associated with Ca^2+^. GSEA analysis and GO enrichment identified Ca^2+^-related DEGs associated with four pathways linked to proteolysis. Since at P13 the *rd1* retina loses mainly rod photoreceptors [12], we then used scRNA-seq to focus on Ca^2+^-related pathways in rods. Also in rods DEGs were enriched in Ca^2+^-related pathways, while, remarkably, this was not the case for cones. The well-studied membrane permeable Ca^2+^ chelator, BAPTA-AM, preserved photoreceptor viability, providing strong evidence for a contribution of excessive Ca^2+^-levels to *rd1* rod degeneration. However, exceedingly low Ca^2+^ levels are also known to trigger cell death [47]. In line with this, BAPTA-AM increased the numbers of dying, TUNEL positive cells in *wt* retina and strongly increased cell death in *rd1*Cngb1^-/-^* double-mutant retina. Together, these findings demonstrate that both exceedingly high Ca^2+^ levels and a depletion of intracellular Ca^2+^ can lead to retinal degeneration [48]. The key question that remains is which Ca^2+^-permeable channel(s) may be responsible for excessive Ca^2+^-influx.

### CNGC and CRAC activity does not promote *rd1* degeneration

In the present study, we investigated the roles of CNGC, CRAC, NCX, and VGCC in photoreceptor cell death (Figure 9). CNGC is activated by high cGMP, linking it to the *rd1* mutation in PDE6. –activated CNGC comprises six homologous members (*CNGA1-4*, *CNGB1* and *CNGB3*), in which CNGA1 and CNGB1 are expressed by rods, while CNGA3 and CNGB3 are expressed in cones [49]. The inhibition of CNGC by L-cis-diltiazem accelerated photoreceptor cell death, in line with previous research [18], but in apparent contradiction with data obtained in *rd1*Cngb1^-/-^* double-mutant mice [17]. However, the *rd1*Cngb1^-/-^* double-mutant mice still retain residual CNGC activity and Ca^2+^ through homomeric CNGA1 channels [50]. L-cis-diltiazem treatment will abolish even this residual Ca^2+^-influx, likely leading to cell death triggered by too low intracellular Ca^2+^.

CRAC activity may also be related to cell death and inhibition of CRAC can increase cell viability [51]. CaMKII potentiates SOCE *via* enhancing Stim1 aggregation and interaction with Orai [32]. In *Pde6b-*mutant mice, the *Camk2g* gene was found to be up-regulated in the present and previous studies [52]. Thus, a contribution of CRAC to Ca^2+^ overload and *rd1* retinal cell death seemed plausible. However, selective CRAC inhibition with CM4620 triggered photoreceptor degeneration in *wt* retina and increased photoreceptor death even further in *rd1* retina, indicating that CRAC-mediated Ca^2+^-influx was important for photoreceptor survival.

NCX is a bi-directional regulator of cytosolic Ca^2+^, capable of mediating both Ca^2+^ influx and Ca^2+^ efflux. NCX inhibition bestows resistance to retinal damage induced by *N*-methyl-D-aspartate (NMDA) and high intraocular pressure [53]. In our hands, the NCX inhibitor SN-6 [54] reduced calpain activity and prolonged *rd1* photoreceptor viability, yet it increased cell death in the *wt* situation. This suggests a reversal of NCX directionality in the two different genotypes, possibly caused by high intracellular Na^+^ levels [31] and membrane depolarization [22]. Thus, cGMP-dependent over-activation of CNGC may have reversed NCX in *rd1* rod photoreceptors. Surprisingly, while BAPTA-AM killed photoreceptors in *rd1*Cngb1^-/-^* retina, perhaps due to Ca^2+^ depletion, NCX inhibition in the *rd1*Cngb1^-/-^* situation attenuated retinal degeneration, indicating that NCX is in forward mode when CNGC activity is low. However, neither BAPTA-AM nor SN-6 preserved the outer retina in the long-term, implying that other mechanisms, such as those triggered by cGMP-dependent protein kinase G (PKG) also promote retinal degeneration [5, 55].

VGCC is activated by membrane depolarization, allowing Ca^2+^ entry into the cells [56]. VGCC consists of four non-covalently associated subunits: α_1_ (L-type and T-type VGCC), β, α_2_δ, and γ [57]. In *rd1* photoreceptors, VGCC may indirectly be activated by cGMP-induced over-activation of CNGC [42]. Hence, D-cis-diltiazem [58], DS5565 [59], and TTA-A2 [60] were used to block different VGCC types. D-cis-diltiazem did not prevent retinal degeneration, consistent with previous literature [61–63]. Furthermore, while we observed an up-regulation of genes coding for the α_2_δ subunits of VGCC in *rd1* rods, treatment with the α_2_δ subunit VGCC ligand DS5565 could not protect *rd1* photoreceptors either. However, DS5565 reduced calpain activity, indicating that the α_2_δ subunit was regulating intracellular Ca^2+^ levels.

Unexpectedly, it was the inhibition of T-type VGCC that significantly attenuated photoreceptor cell death. This is surprising since T-type channels are generally thought to open at relatively negative membrane potentials of around –40 to –80 mV, as opposed to L-type channels which may open already at –20 mV [64]. In photoreceptors the resting potential is approx. –40 mV and the membrane potential would reach –70 mV only during light-induced photoreceptor hyperpolarisation [7]. Thus, in *rd1* photoreceptors, which cannot be hyperpolarized by light due to *PDE6* dysfunction, one would assume that L-type VGCC rather than T-type should carry most of the Ca^2+^-currents. However, we cannot exclude the possibility that T-type channels change their properties to become permeable at less negative membrane potential, or that *rd1* photoreceptor membrane potential shifts to more negative values.

Overall, our data suggest T-type VGCC and NCX contribute to *rd1* retinal degeneration. One may speculate that in the initial phases of an individual photoreceptoŕs demise, the membrane potential is still maintained at sufficiently negative values, allowing for an opening of T-type channels and Ca^2+^-influx. Eventually, however, the cell may no longer be able to keep its membrane potential and becomes depolarized, so much so that NCX reverses its direction and turns into a net importer of Ca^2+^. This idea of two different stages in the dysregulation of photoreceptor Ca^2+^-levels may also help to understand why NCX inhibition did not afford long-term protection. Whatever the case, our data highlights T-type VGCC as a potential target for therapeutic intervention.

### Calpain-2 causes photoreceptor cell death and negatively regulates calpain-1

High levels of intracellular Ca^2+^ will activate calpain-type proteases and calpain activation may be connected to CNGC activity [14]. Calpain consist of a family of Ca^2+^-activated neutral cysteine proteinase involved in a large number of cellular processes [37, 65]. Originally, the calpain proteolytic system was reported to comprise three different proteins: calpain-1 (or μ-calpain, activated at μM Ca^2+^ concentrations) and calpain-2 (or m-calpain, activated at mM Ca^2+^ concentrations), as well as their endogenous inhibitor, calpastatin [65]. Recent evidence indicates that calpain-1 and calpain-2 play opposite roles, with calpain-1 being neuroprotective while calpain-2 promotes neurodegeneration [37]. We previously found that in *rd1* mice, activated calpain-2 was increased rather than calpain-1 [36], and the current GSEA analysis indicated a positive regulation of proteolytic pathways. Thus, we assumed that calpain-2, as part of the proteolytic processes induced by Ca^2+^ [66], led to retinal degeneration. Therefore, we treated *rd1* retina with the calpain-2 specific inhibitor NA-184 [38]. This caused a significant decrease in the numbers of TUNEL positive cells in the ONL, highlighting the role of calpain-2 in photoreceptor cell death. On the other hand, calpain-1 activation was decreased in situations where calpain-2 activation was high, possibly due to calpain-2-dependent cleavage of calpain-1’s protease core domain 1 [67]. Accordingly, NA-184 treatment also increased calpain-1 activation. These findings extend one of our previous studies where we found the general calpain inhibitor calpastatin peptide to protect photoreceptors *in vitro* and *in vivo* [68]. Yet, since calpastatin peptide inhibits both calpain-1 and –2, it seems likely that the beneficial effects of calpain-2 inhibition were partly offset by the additional calpain-1 inhibition. Taken together, we found calpain-2 activation to be related to photoreceptor cell death, and this was associated with decreased calpain-1 activation.

### Ca^2+^ differentially regulates PARP and sirtuin activity

PARP and HDAC activity were previously connected to *rd1* degeneration and to Ca^2+^-signalling [14, 69, 70]. PARP is a DNA repair enzyme, which catalyses the polymerization of ADP-ribose units – derived from the ADP donor NAD^+^ – resulting in the attachment of either linear or branched PAR polymers to itself or other target proteins [71]. However, excessive PARP activity may drive a specific form of cell death, termed PARthanatos [72], which has also been connected to in retinal degeneration [4]. Accordingly, in certain IRD animal models, PARP inhibition was found to be neuroprotective [14, 73]. In line with previous literature [74, 75], the inhibition of Ca^2+^ –permeable channels reduced PARP activity and PAR generation in *rd1* explants, suggesting that Ca^2+^ regulated PARP activity. However, it is unclear how Ca^2+^ produces PARP over-activation. The specific calpain-2 inhibitor, NA-184, did not reduce PARP activity in the *rd1* explants, suggesting that Ca^2+^ controls PARP activity independently of calpain activity [14].

HDACs catalyse the removal of acetyl groups from lysine residues of both histone and nonhistone proteins [76]. HDACs are divided into zinc-dependent HDACs, and sirtuins, a family of NAD^+^-dependent HDACs [76]. As PARP is a main consumer of NAD^+^ [77], sirtuin activity can be regulated by PARP according to the NAD^+^ levels [78]. The Sirtuin-1 protein is suggested to be protective in neurons [79]. While we observed decreased PARP activity after most of our experimental interventions, sirtuin activity was increased after treatment with BAPTA-AM and the NCX inhibitor SN-6, suggesting that the resultant decrease in PARP activity may relieve the NAD^+^ shortage induced by PARP. Interestingly, calpain-2 inhibition also increased sirtuin activity without affecting PARP activity. However, currently there is no evidence indicating that calpain-2 cleaves HDACs. Still, HDACs may regulate calpain-2 activation indirectly *via* epigenetically increasing calpastatin expression [80]. Calpain may also influence mitochondrial biogenesis, as calpain plays a detrimental role upstream of the peroxisome proliferator-activated receptor γ (PPARγ) coactivator-1α (PGC-1α) pathway [81], which regulates the transcription of numerous nuclear-encoded mitochondrial genes [82] and is a key driver of mitochondrial biogenesis [83]. Thus, it is possible that calpain-2 inhibition improves *rd1* photoreceptor energy metabolism.

## CONCLUSION

In IRD research, the role of Ca^2+^-signalling in disease pathogenesis has for a long time remained controversial. Our results now demonstrate that Ca^2+^-signalling can have both beneficial and detrimental effects in *rd1* photoreceptors, depending on the source of Ca^2+^ and perhaps its intracellular localization. More specifically, inhibition of CNGC and CRAC accelerated cell death, while Ca^2+^ chelation, as well as inhibition of NCX and T-type VGCC increased photoreceptor viability. Likewise selective inhibition of calpain-2 improved photoreceptor viability. Remarkably, activation of calpain-1 and sirtuin-type HDAC was linked to photoreceptor survival. In contrast, general calpain activity and activity of PARP was found to be destructive. While our results propose T-type VGCC and calpain-2 as attractive targets for IRD therapies (Figure 9), a careful context– and genotype-specific evaluation seems warranted as shown, for instance, by the opposing results obtained in *rd1* single-mutant and *rd1*Cngb1^-/-^* double-mutant retina. Overall, our study illustrates the complexity of Ca^2+^-signalling during photoreceptor degeneration and highlights the need to clearly delineate destructive and protective pathways to allow for the rational development of new therapeutic approaches for IRD and related retinal diseases.

## ACKNOWLEDGEMENTS

The authors would like to thank Michel Baudry (CDM, Western University of Health Sciences, Pomona, CA) for providing NA-184, and editing the manuscript. We thank Norman Rieger (from Institute for Ophthalmic Research, Eberhard-Karls-Universität Tübingen) for excellent technical assistance.

## AUTHOR CONTRIBUTIONS

Conceptualization, J.Y. and F.P.-D.; methodology, J.Y., L.W., X.H., Y.D., K.J. and Z.H.; software, Q-L.Y., Q-X.Y. and J.Y.; validation, J.Y. and L.W.; formal analysis, Q-L.Y. and Q-X.Y.; investigation, J.Y. and L.W; data curation, J.Y.; writing-original draft preparation, J.Y., L.W. and Q-L.Y.; writing—review and editing, F.P-D.; visualization, J.Y. and Q-L.Y.; supervision, F.P-D.; project administration, F.P-D.; funding acquisition, F.P-D. and Z.H All authors have read and agreed to the published version of the manuscript.

---

**Figure S1.**
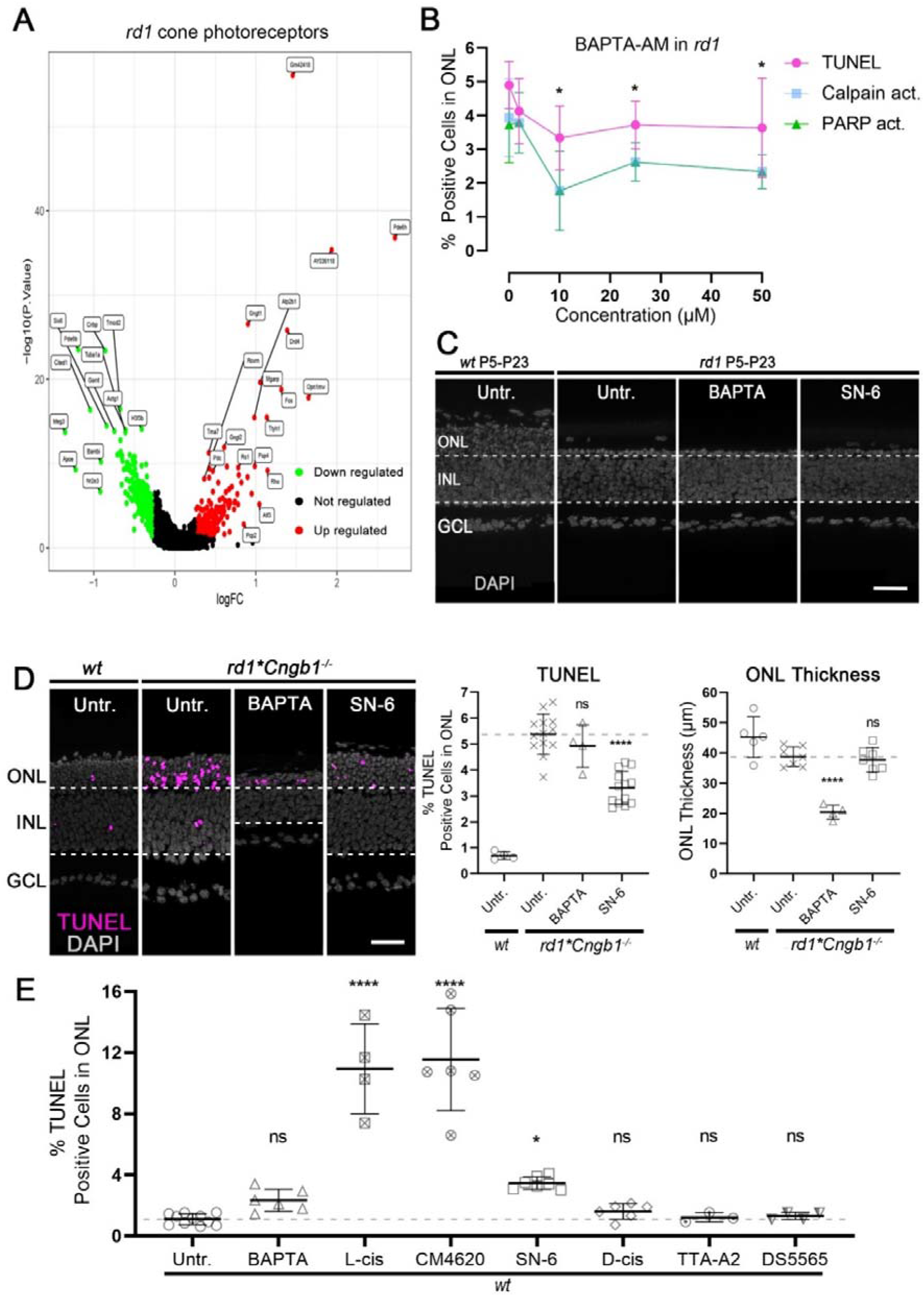
Bioinformatic analysis and effects of interventions targeting photoreceptor Ca^2+^-permeable channels. **A**) Differentially expressed genes (DEGs) in *rd1* cone photoreceptors at post-natal day (P)13. **B**) Dose-response curve for BAPTA-AM in *rd1* explant cultures. In the outer nuclear layer (ONL), 10, 25, and 50 µM BAPTA-AM significantly reduced calpain activity, PARP activity, and cell death as detected via the TUNEL assay. **C**) Long-term treatment from P5 to P23, with 10 µM BAPTA and 40 µM SN-6, in *rd1* explant cultures, compared to untreated (Untr.) wild-type (*wt*) and *rd1* specimens. **D**) Effect of 10 µM BAPTA-AM and 40 µM SN-6 treatments on *rd1*Cngb1^-/-^*retinal cultures. TUNEL and photoreceptor thickness in treated retinas compared to Untr. *rd1* specimens. **E**) Different interventions targeting Ca^2+^-permeable channels in *wt* retinal explant cultures. The scatter plots show the percentage of TUNEL positive cells in the ONL. Scale bars in C, D. 50 µM. Statistical significance was assessed using one-way ANOVA and Tukey’s multiple comparison post hoc test. Untr.: n=8 retinal explants from different animals; BAPTA: 5; L-cis: 4; CM4620: 6; SN-6: 7; D-cis: 6; TTA-A2: 3; DS5565: 4.

**Figure S2.**
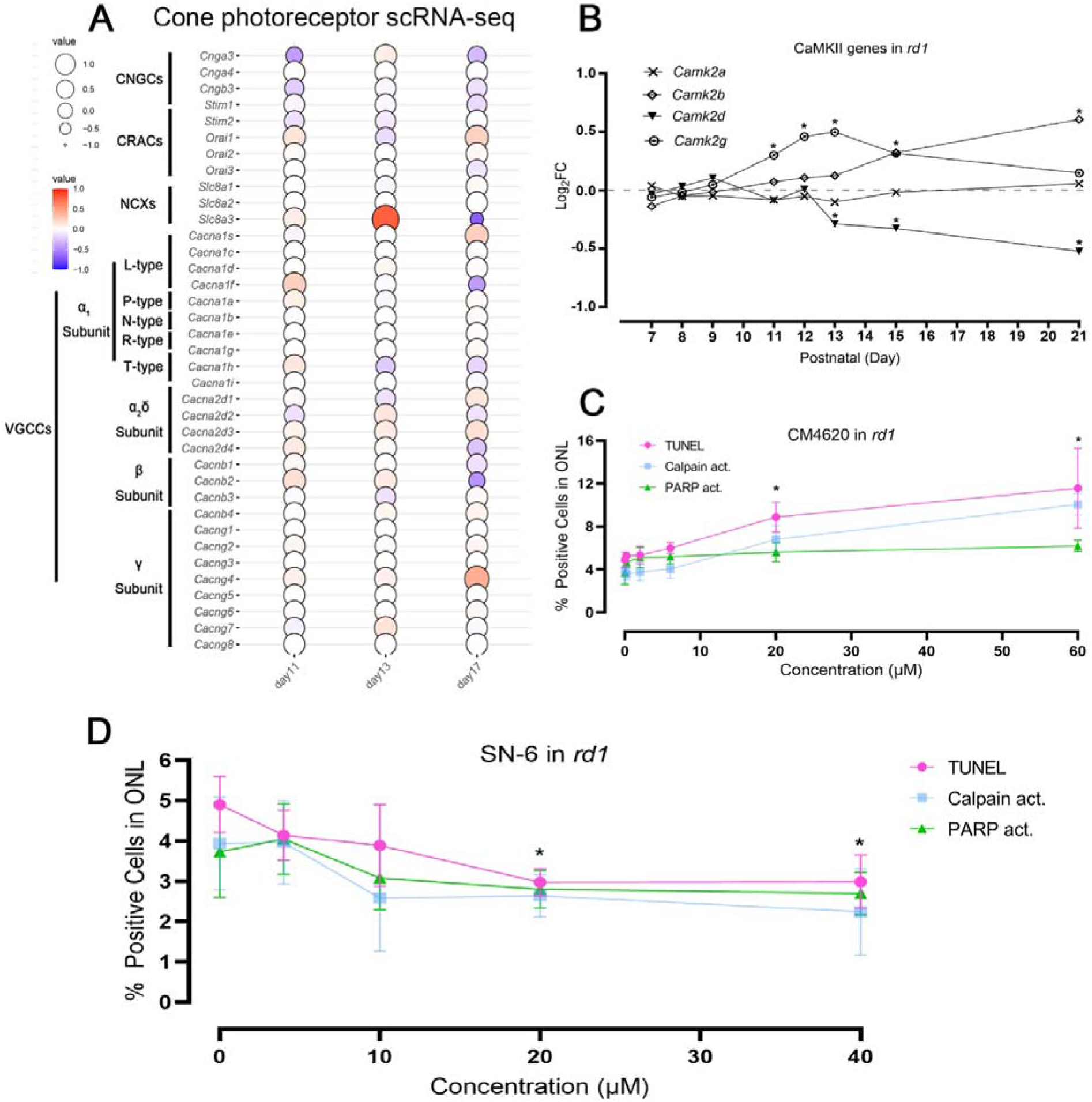
Expression of *Camk2* genes and dose-response curves for CM4620 and SN-6. **A**) Balloon plot showing time-dependent expression changes (post-natal day (P) 11 to P17) of cyclic nucleotide–gated channel (CNGC), Ca^2+^-release activated channel (CRAC), Na^+^/Ca^2+^ exchanger (NCX), and voltage-gated Ca^2+^ channel (VGCC) in *rd1* cone photoreceptors. **B**) Analysis of the expression of Ca^2+^/calmodulin-dependent protein kinase II (CaMK2) genes during *rd1* photoreceptor degeneration. **C**) Different concentrations of CM4620 were tested in *rd1* explant cultures. In the outer nuclear layer (ONL), at concentrations of 20 µM and 60 µM, CM4620 significantly increased ONL calpain activity, PARP activity, and cell death, as assessed by the TUNEL assay. **D**) Dose-response for SN-6 in *rd1* explant cultures. 20 µM and 40 µM SN-6 significantly reduced calpain activity, PARP activity, and cell death (TUNEL) in the ONL. Statistical significance was assessed using one-way ANOVA and Tukey’s multiple comparison post hoc test.

**Figure S3.**
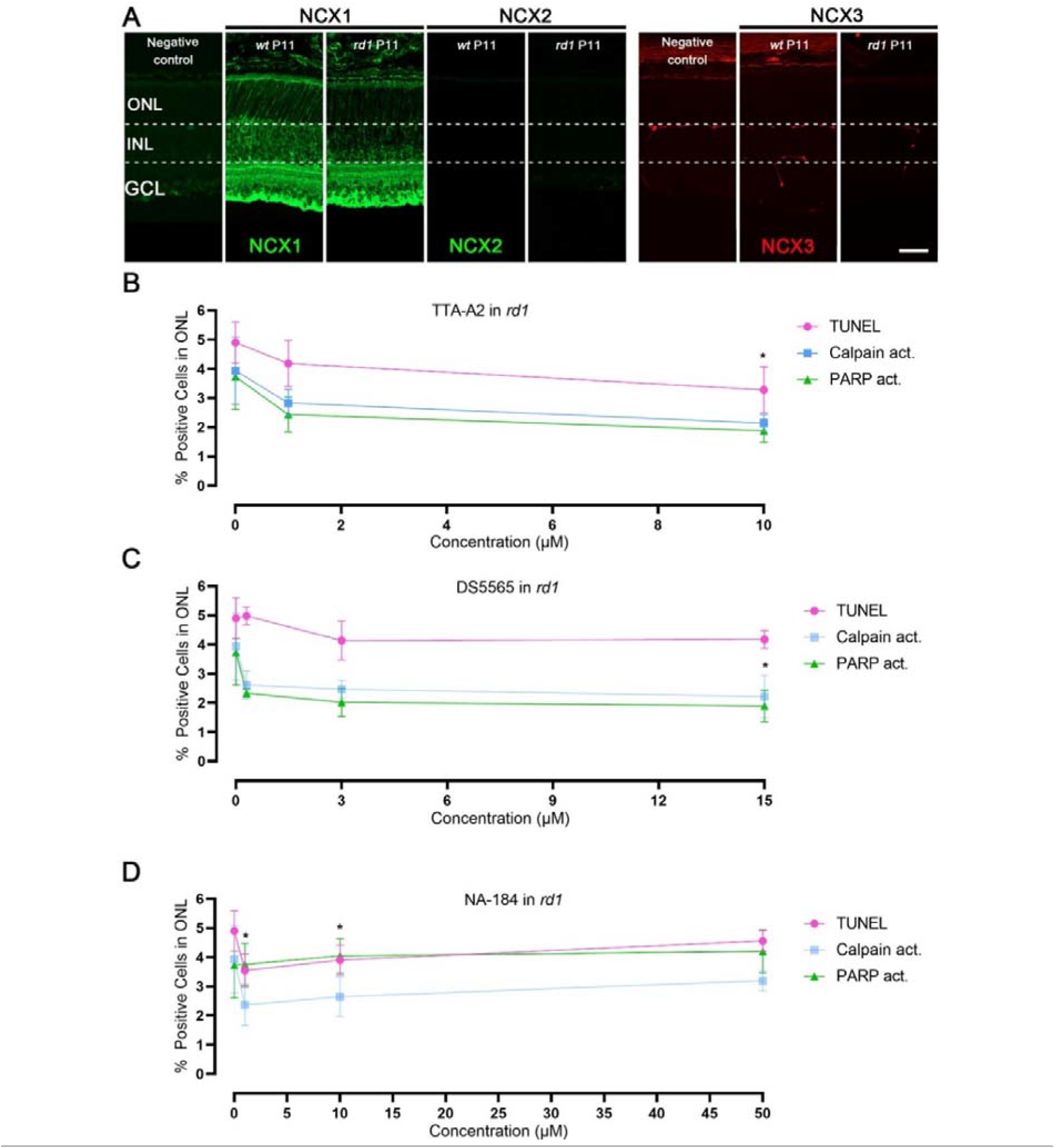
Retinal NCX expression and dose-response curves for TTA-A2, DS5565, and NA-184. **A**) Immunostaining of Na^+^/Ca^2+^ exchanger (NCX) family. NCX1 was found to be expressed in both inner and outer retina, while NCX2 and NCX3 were not detected. **B**) Different concentrations of TTA-A2 were tested in *rd1* explant cultures. In the outer nuclear layer (ONL), 10 µM TTA-A2 significantly reduced calpain activity, PARP activity, and cell death as assessed with the TUNEL assay. **C**) Dose-response for DS5565 in *rd1* explant cultures. 15 µM DS5565 significantly reduced calpain activity, PARP activity, but not cell death (TUNEL) in the ONL. **D**) Dose-response for NA-184 in *rd1* explant cultures. At concentrations of 1 µM and 10 µM NA-184 significantly reduced ONL calpain activity and cell death (TUNEL) but did not decrease PARP activity. Statistical significance was assessed using one-way ANOVA and Tukey’s multiple comparison post hoc test.

**Supplemental Table 1.**
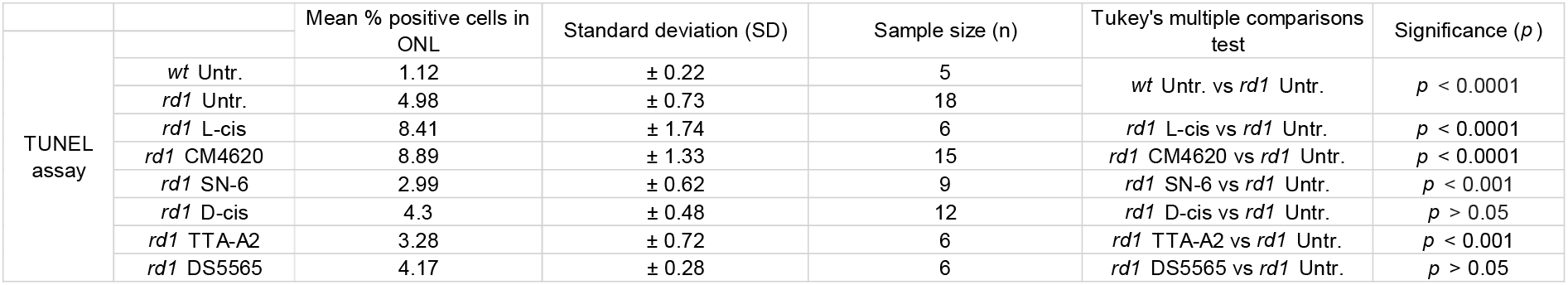
– Quantification of TUNEL positive, dying cells in the outer nuclear layer (ONL).

**Supplemental Table 2.**
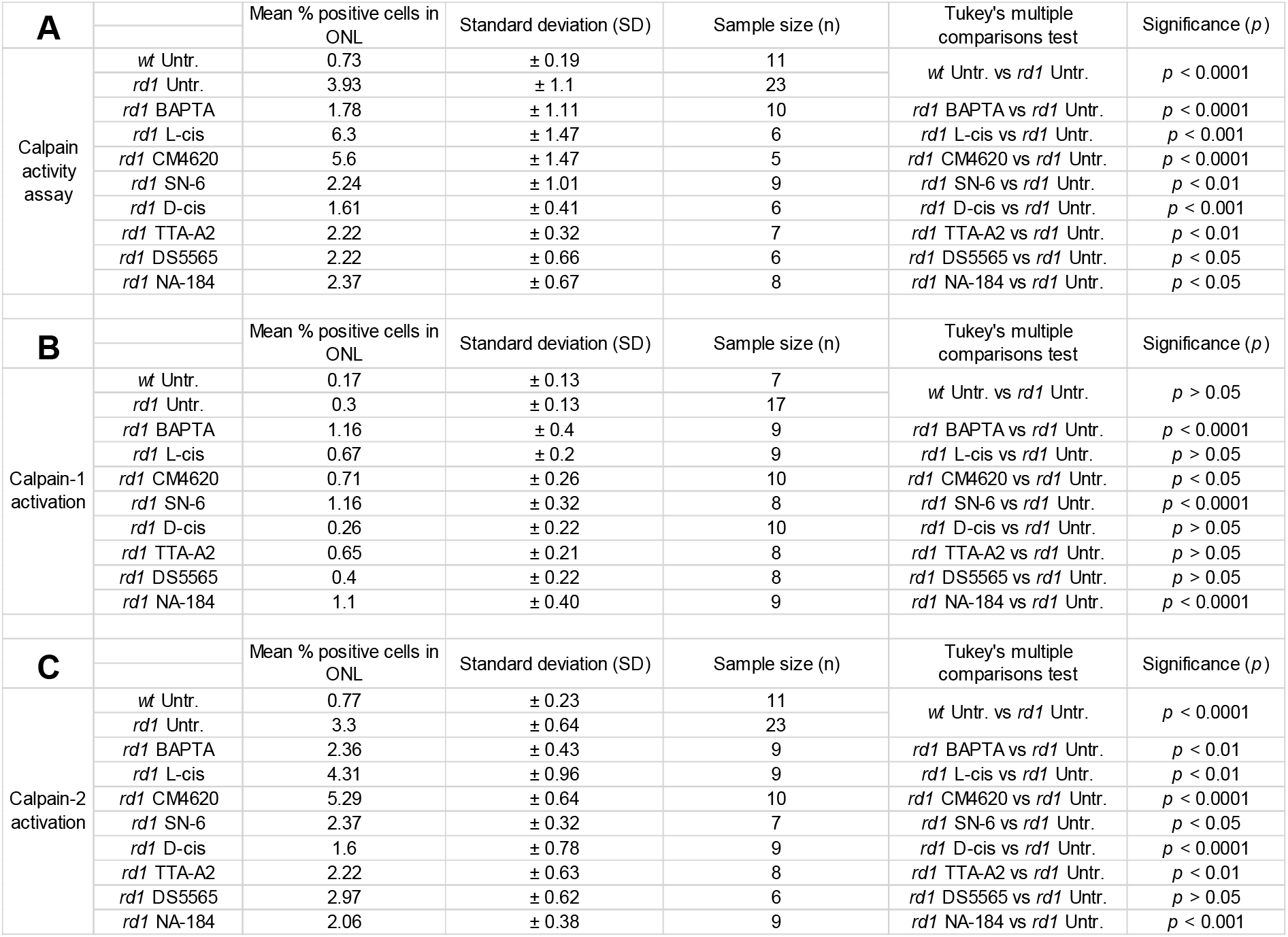
– Quantification of cells positive for calpain activity/activation in the outer nuclear layer (ONL).

**Supplemental Table 3.**
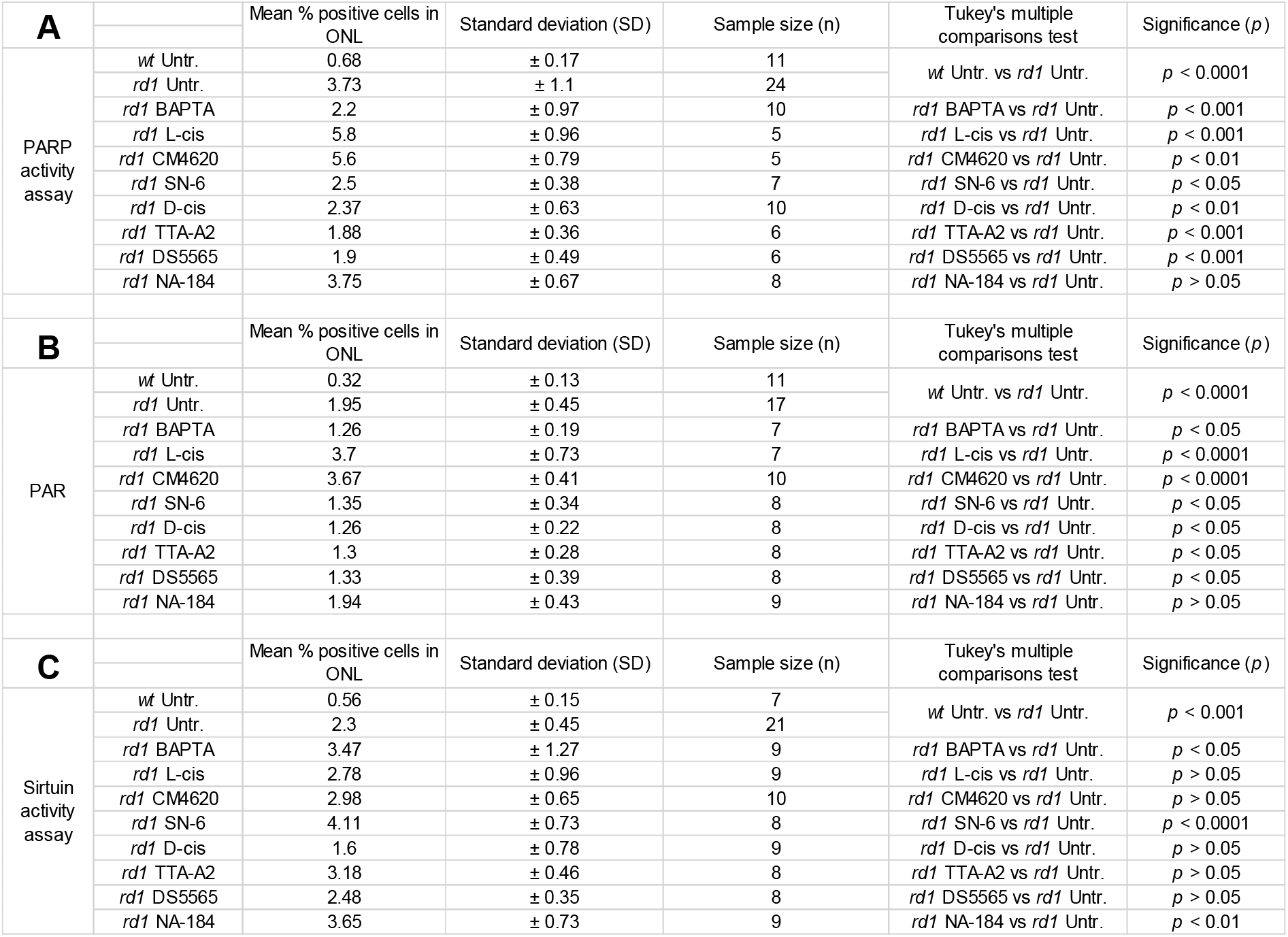
– Quantification of cells positive for PARP activity, PAR accumulation, and Sirtuin activity in the outer nuclear layer (ONL).

