## Supplementary figures and images for "Inherited retinal degeneration: T-type voltage-gated channels, Na^+^/Ca^2+^-exchanger and calpain-2 promote photoreceptor cell death"

### Supplemental figure 1

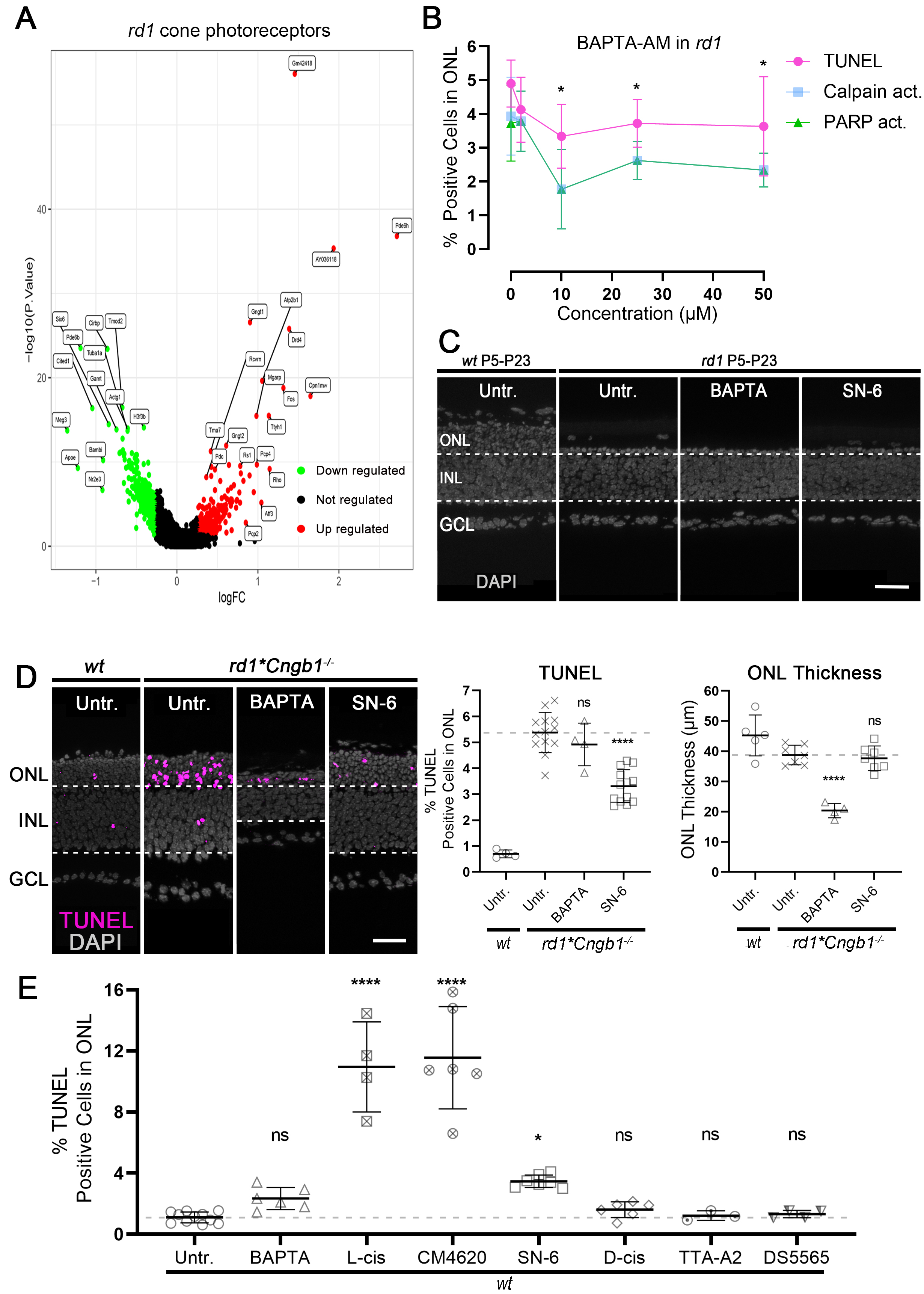

### Supplemental figure 2

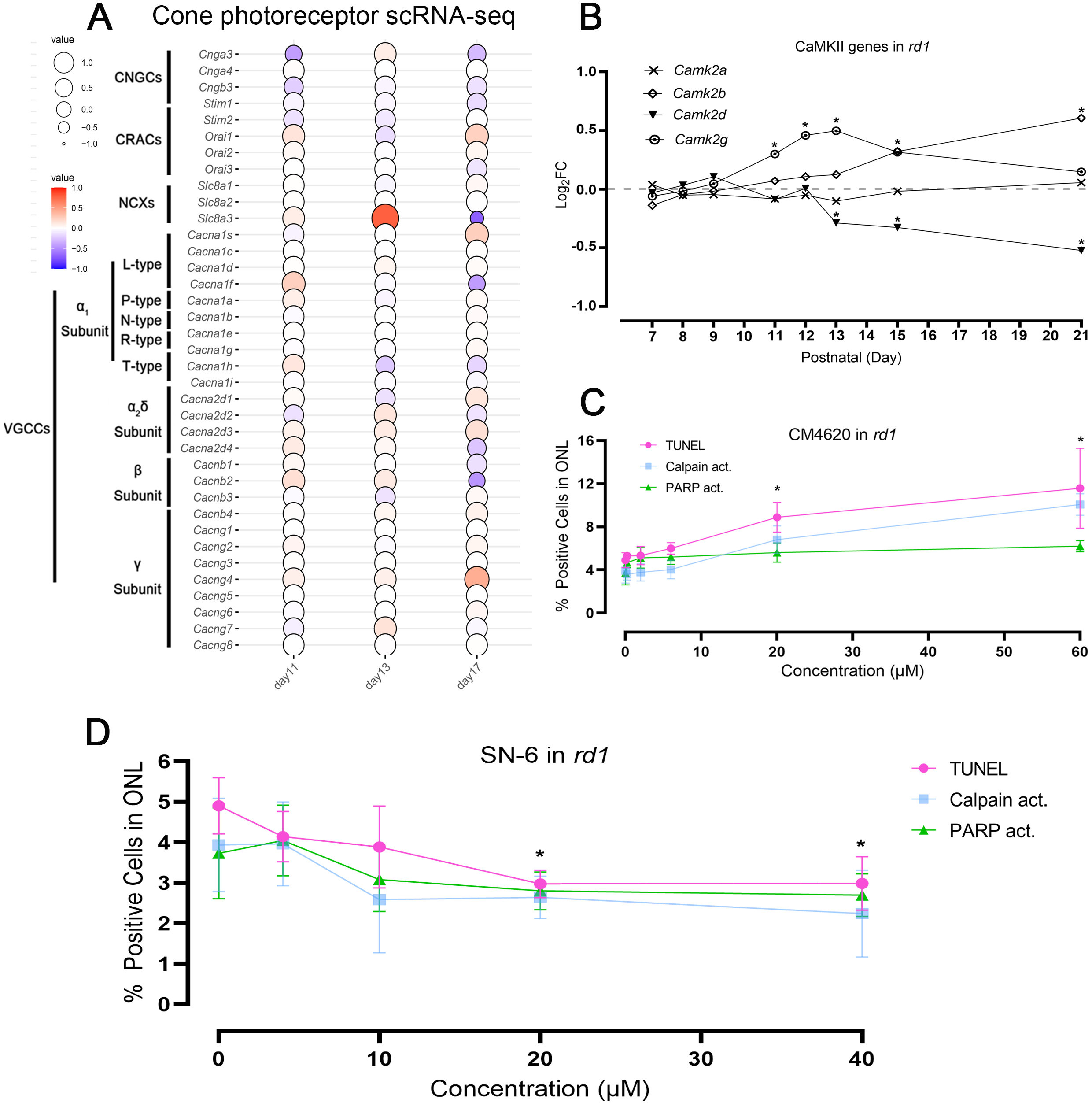

### Supplemental figure 3

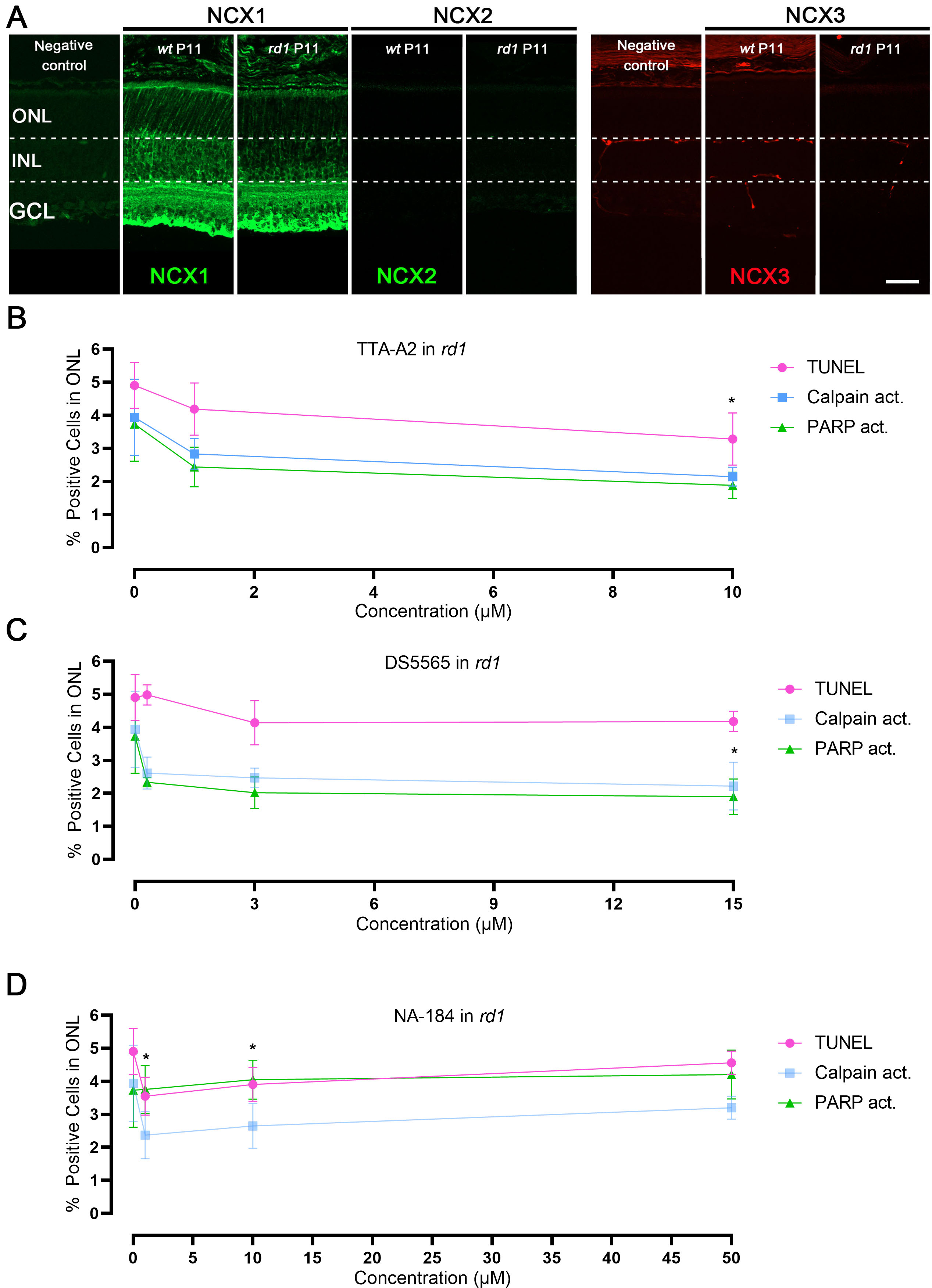

### Supplemental table 1

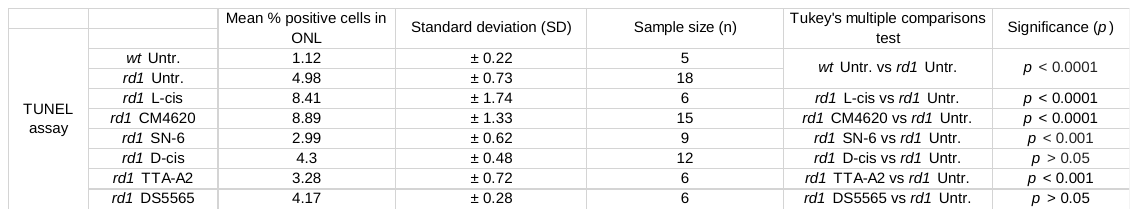

### Supplemental table 2

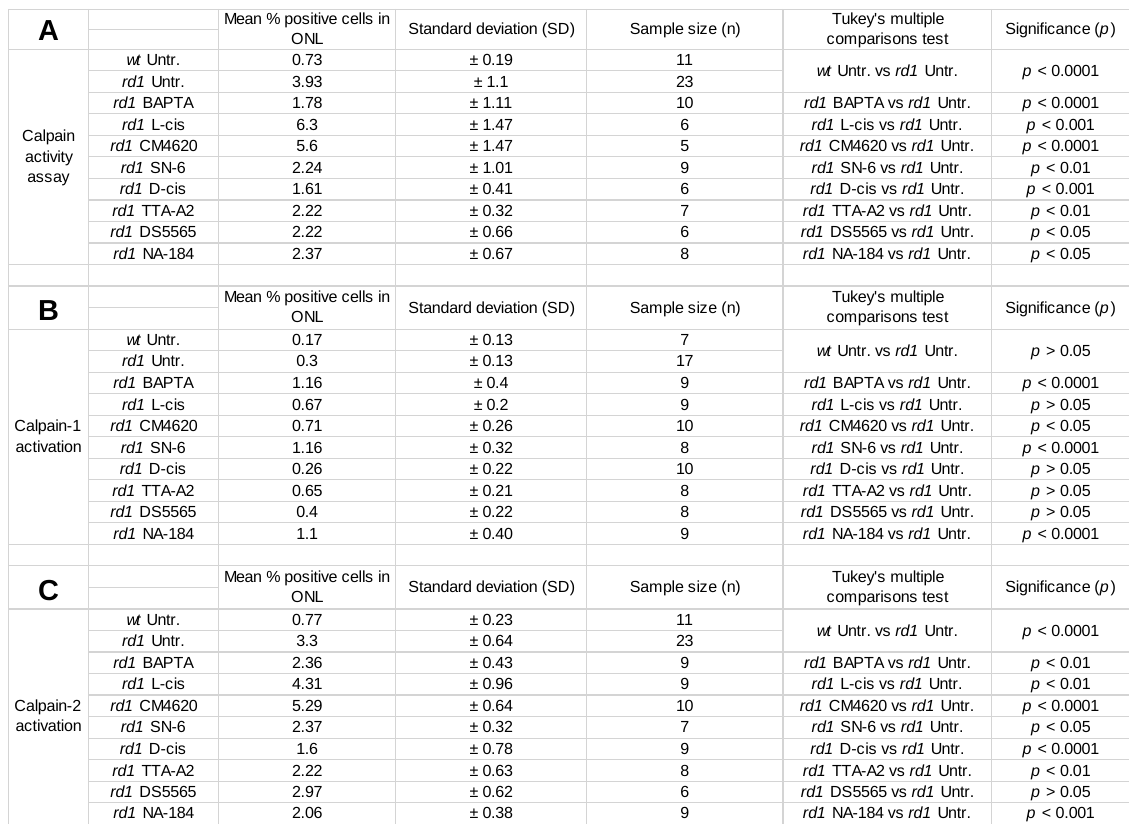

### Supplemental table 3

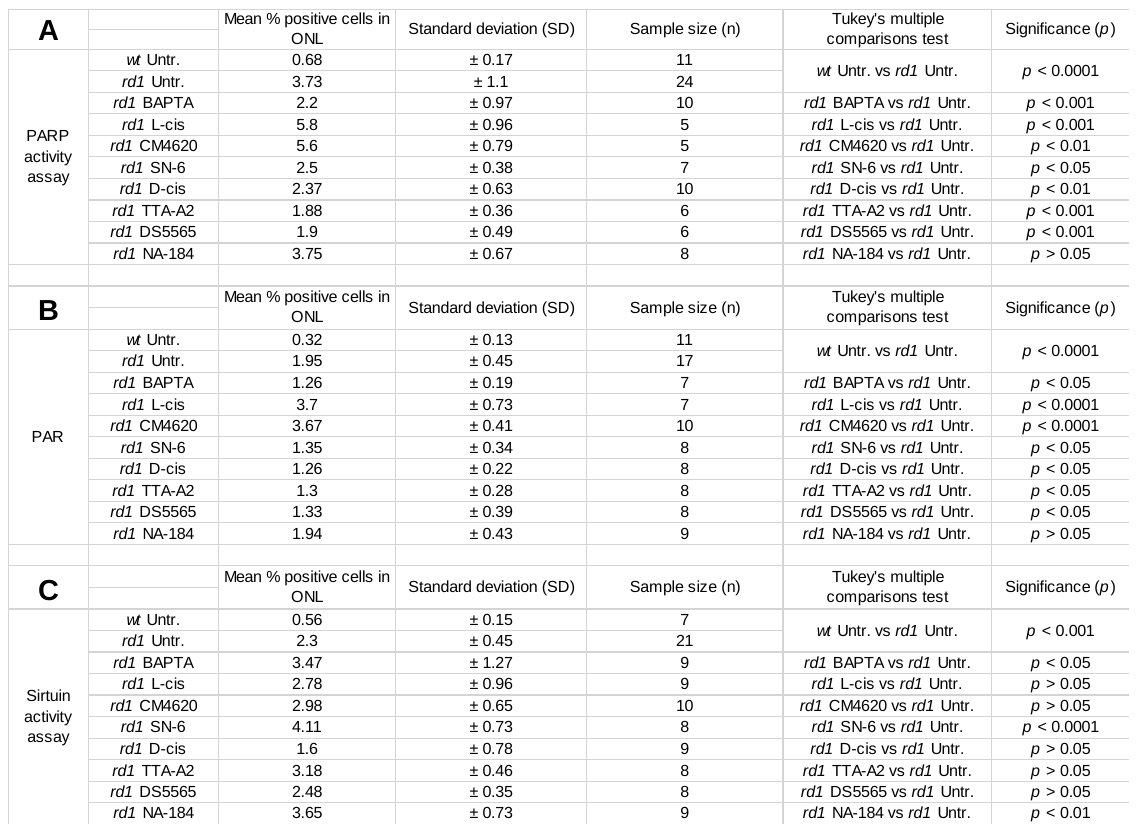
